# Quantitative modelling of biological response dynamics reveals novel patterns in plant volatile signalling

**DOI:** 10.1101/2025.11.26.690448

**Authors:** Jamie M. Waterman, Gareth J. Moore, Loren K. Amdahl-Culleton, Sara Hoefer, Matthias Erb

**Author notes:** Authors contributed equally to this work (Jamie M. Waterman, Gareth J. Moore).

## Abstract

Biological responses to environmental stimuli are inherently dynamic. Recent technological advances enable detailed time-resolved measurements of such responses across biological systems. However, a consistent, universal standard for quantifying temporal response dynamics is lacking, thus limiting biological insights and comparisons. We developed an unbiased mathematical model structure that allows for the quantification of biological response dynamics without *a priori* knowledge of underlying biochemical mechanisms. Using the model to quantify the dynamics of stress-induced plant volatiles, we uncover a range of novel patterns in volatile signalling, including i) a strong light-independent impact of the time of day of wounding on the onset, duration and shape of the volatile induction responses, ii) an accentuation of volatile-specific induction curve shapes by herbivory-associated molecular patterns (HAMPs) and iii) independent regulation of the strength and duration of volatile induction across genotypes. We also demonstrate that our approach performs well across diverse plant and animal systems, potentially indicating universality across responses. The model is also robust to partial response curves, low resolution data and complex multi-modal responses arising from overlapping stimuli, enabling identification of priming events from otherwise convoluted curves. As all responses measured conform to a common model structure, yet parameter values diverge markedly, we conclude that biologically meaningful information is ignored when temporal dynamics of responses are not quantified. The presented approach will pave the way to identifying new response patterns, and their function, across the tree of life.

## Introduction

Inducible biological responses are transient in nature, changing substantially over relatively short periods of time (*1–5*). By consequence, real-time measurements are crucial to understand physiological processes, precisely diagnose chronic and acute stress patterns and resolve the mechanisms behind ecological phenomena (*6–11*). Although these responses are inherently dynamic, quantitatively describing and comparing their temporal features remains a challenge. It is well established that static features such as mean concentration of an analyte at a given instance are useful for general interpretations of stimulus-response relationships, such as the inverse relationship between viral load and effective immune response in humans and the positive relationship between herbivore damage and volatile emissions in plants (*12*, *13*). However, static values, even if measured at different time points, ignore meaningful, quantifiable features of the dynamics themselves.

Recent examples illustrate the potential of moving beyond measures such as static means to generate new biological insights relevant to inducible responses. In plants, variation in the spatial distribution of a toxic metabolite, either within a plant or between plants, can affect herbivore feeding behaviour and thus fitness parameters such as growth, independently from the total concentration of said toxin (*14–16*). In animals, dendritic signals are sensitive to temporal patterns of synaptic activation; irrespective of the amount of signal, the temporal patterns of signal perception shape spike outputs, which drive crucial functions such as sound localisation (*17*, *18*). Additionally, the interaction between vaccination status, viral exposure and immune responses are all temporally linked, and the effects/response of each will be dependent on how the others change over time (*19*).

From a mechanistic perspective, the dynamics of inducible responses are determined by the interplay of highly regulated signalling events (*20–25*) and modifications to one or more events can have far-reaching consequences on response dynamics and metabolic outcomes (*1*, *17*, *26–28*). For diffusion-based processes such as generation of reactive oxygen species (ROS) or single-cell signalling events like MAPK cascades, the underlying steps are few and well-characterised, and relatively simple, real-time mechanistic models have been developed (*6*, *29–32*). Some models of temporal dynamics of metabolism with more complex biosynthetic pathways have also been developed (*33*, *34*). These models are valuable for elucidating biochemical mechanisms and understanding the complexity of metabolic regulation. However, they require substantial *a priori* biochemical knowledge and the ability to measure numerous kinetic and regulatory parameters (steps), making their construction challenging even for well-characterised metabolic processes (*35–37*). For more complex and less-well characterised pathways, such as the *de novo* production of specialised metabolites, complete time-resolved mechanistic models are not currently feasible. Many underlying processes, including enzyme kinetics, precursor and product concentrations, transport delays, degradation pathways and other regulatory processes remain unknown (*38–40*). Because the biochemistry is not fully defined, the number and nature of parameters required to describe such processes are effectively arbitrary, leading to under-constrained models in which multiple parameter combinations can reproduce the same behaviour and limit reliable fitting and meaningful interpretation (*37*). Moreover, existing models are often system-specific and must be adapted depending on the metabolites or pathways studied. Other existing approaches, such as template mode of variation (TMV), are useful for parameterising response curves, but parameterisations are relative to a mean template defined for a pre-defined set of curves, meaning that the resulting parameter values depend on the set of response curves included in the analysis (*41*, *42*). Consequently, there is no universally standardised framework or parameterisation that can be easily implemented across systems, which limits comparability, broad adoption and ultimately accessibility of dynamics quantification.

Mechanistically informed models may not always be necessary to study biological responses. Volatile chemical signals emitted by organisms shape important interactions such as those between insects and their hosts (*43*, *44*). In this case, the mechanism of volatile formation matters less than when this chemical information is released, how long it persists and whether amounts produced cross perceptual thresholds (*10*, *45–47*). These characteristics are also important for within-organism biochemical processes. Consider protein dynamics (i.e., moving between different conformational states over time), for which similar dynamic features would determine interactions with other proteins and ligands, and thus shape the fate of biochemical processes (*48*, *49*). Thus, extracting time-resolved features of response dynamics that are intuitive, broadly comparable and important for driving biological functions becomes important.

Motivated by the need to extract biologically meaningful information from dynamic responses, and the lack of a robust framework to do so, we set out to develop a method that allows for biologically meaningful, consistent and interoperable comparisons of dynamic responses without explicit knowledge of the underlying physiochemical and biochemical processes. We used our model quantify the dynamics of diverse plant volatile responses to different stimuli to identify and evaluate whether meaningful differences exist in the dynamics themselves, generating novel biological insights. We considered plant responses to stress to be an optimal test case, as plants have evolved a highly complex array of responses to environmental stimuli, that cover a broad range of physiological and ecological functions (*50*), and show starkly distinct kinetics (*44*). Induced volatiles in particular can be measured non-destructively, enabling diverse processes to be observed in real time (*7*). Finally, these volatiles provide important chemical information to the wider environment, and we thus assume that the generated insights will be ecologically informative (*10*, *47*). Further, we also test our approach on published data from other signalling dynamics across plant and animal species (*51–56*).

Through this approach, we uncovered that all measured responses conform to a common model structure, while the parameter values varied considerably between stimuli and response types. Not only would these phenomena go undetected without a robust quantitative framework, but they reveal that biologically meaningful information is encoded in induced response dynamics and that temporal response structure carries functional significance. Importantly, the model structure is not specific to either response type or timeframe, and is explicitly applicable over a broad range of dynamic processes and biological systems, thus unlocking novel comparative and integrative approaches across the tree of life.

## Results

### Model development

To build a phenomenological model that captures dynamic responses in an unbiased manner, we modelled responses as a distribution of waiting times. We asked the question: *How long does it take for a response unit (e.g. a volatile molecule) to appear following a stress event?* In doing so, we interpret each measured unit as a probabilistic occurrence with a certain delay, and thus model the response curve as a probability density function.

This framing is advantageous for three reasons: first, the model yields a relatively small number of parameters that are both identifiable (i.e. are intuitive by eye) and interpretable (i.e. have biological meaning). This minimises overfitting and, as a result, makes the model robust to sample-to-sample variation. Secondly, it enables comparability across responses which involve very different mechanisms and timeframes. Thirdly, the same framework can be applied to any induced phenotype that can be measured quantitatively over time, such as early signalling events, gene expression, hormone accumulation, enzymatic conversions, volatile emissions and resistance induction, even if they show starkly different dynamics and biochemical processes, as they all typically exhibit the same behaviour: induction delay following stimulation and a rapid increase in response, which, after reaching a peak, gradually declines (*1*, *2*, *28*, *57–65*). This is mathematically intuitive as these responses are built from a sequential chain (or combinatory network) of activation, synthesis, transport and eventually measurement/detection. Each step adds stochastic waiting time with their sum naturally producing a skewed, unimodal distribution (*66*). Modelling this kind of process usually involves treating each biochemical step as a stochastic waiting time drawn from a defined probability distribution and then combining these into a final waiting-time distribution which captures the total observed response dynamics. As such, we used a reparametrised gamma distribution, which retains its form when steps are combined and can thus describe both early and later phases of the response in a consistent way (see materials and methods for full model details):

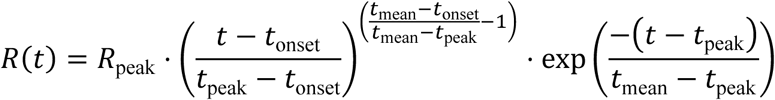

where:

• *R_peak_* is the maximum measured response,

• *t_onset_* is the onset delay,

• *t_peak_* is the time of peak response,

• *t_mean_* is the mean response time.

The fitting parameters can then be used to calculate additional characteristic parameters (Fig 1A).

**Figure 1.**
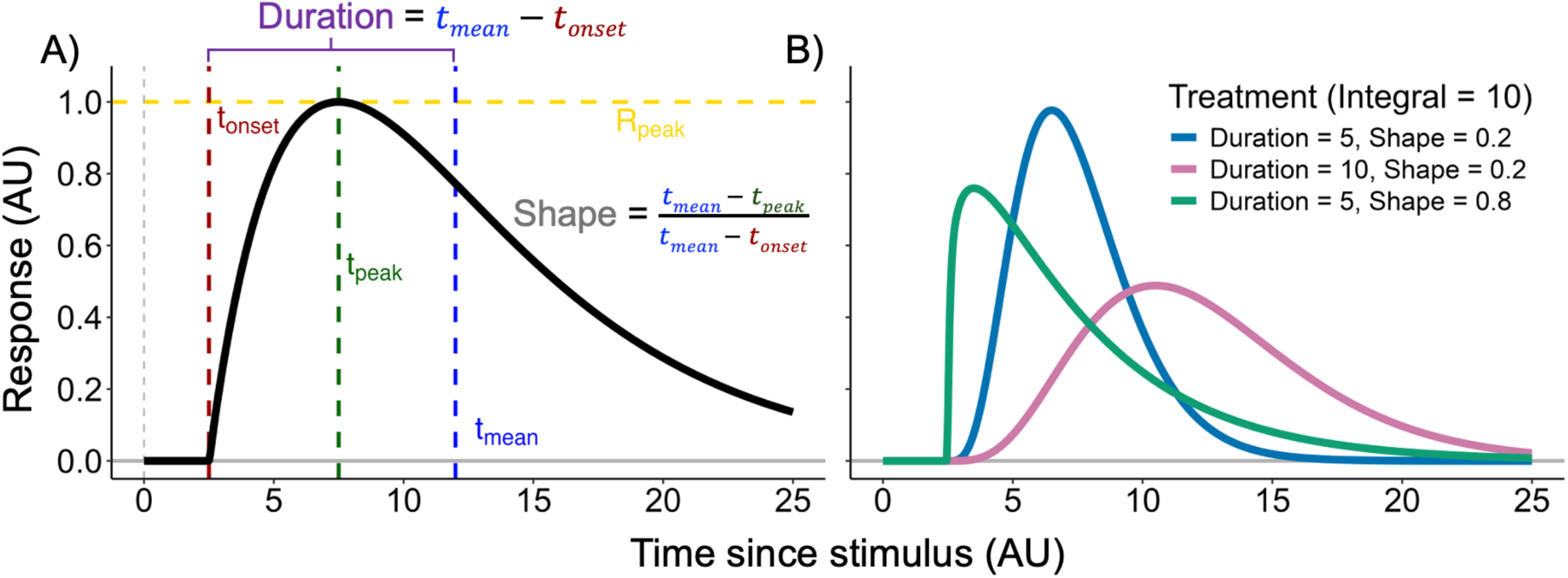
Theoretical examination of important features of biological response curves. A) Parameters of the model include *t_onset_*: onset delay, *t_peak_*_’(_: time of peak response, *t_mean_*: mean response time, *R_peak_*_’(_: the maximum measured response. These parameters are further reparametrised into *Duration*: metric of the length of response and *Shape*: our symmetry feature, which for gamma-like distributions falls between zero and one. B) *In silico* data highlighting the utility of model parameters, namely that discrete temporal response patterns can emerge, despite identical integrals.

Total response (*Integral*) – The integral is often difficult to measure experimentally; due to time limitations in the measurement processes, entire curves are not resolved and thus incomplete integrations are used (*67*). When fit onto the experimental data, our model can be used to calculate the theoretical integral of the response curve even if it is incomplete:

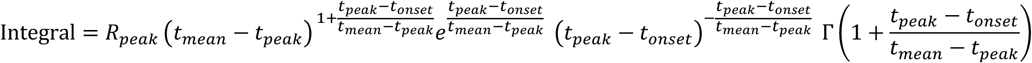

where Γ() is the Gamma function, a continuous generalisation of the factorial (*46*, *47*).

#### Duration

For response dynamics, response length is typically defined as ‘broadness’, for example as full width at half max (FWHM) (*6*). The FWHM of the Gamma function has no analytical solution and must be determined numerically (*70*). Additionally, FWHM, does not explicitly capture the start of the response. For these reasons *Duration* is used as a simple alternative, achieving a similar descriptive value and used as a normalisation value in the ‘shape’ feature below.

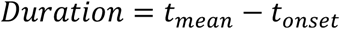

#### Shape

The ‘shape’ of a response curve is an abstract feature. Since our model is intended to be used over variable time scales, we developed a time-normalised (by *Duration*) metric that allows for comparison of curve shapes independently from response length. Further, we aimed to understand the symmetry of the curve in response to different stimuli or across plant genotypes/species to broadly assess temporal nuances. *Shape* presents a new phenotypic axis which may link to important physical of biochemical properties that regulate induced responses. *Shape* is a normalised value, and when responses follow gamma distributions it falls between zero and one; a *Shape* of zero indicates a perfectly symmetrical curve (*t_peak_* = *t_mean_*) and a shape of one represents a completely right-skewed curve (*t_peak_* = *t_onset_*):

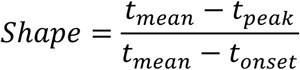

Taken together, the fit and derived parameters give a framework for comparing curves, particularly those that appear distinct by eye but are difficult to distinguish quantitatively (Fig 1B).

The model described the observed responses well, with a mean R² of 0.815 and a normalised RMSE of 0.129 across individual fitted curves. Fit quality was consistently higher when mean fitted curves were compared with mean observed curves within treatment groups (R² = 0.942, normalised RMSE = 0.066), indicating that most of the residual variation at the individual level reflects replicate-to-replicate noise rather than a failure of the model to capture the underlying response shape (Supplemental Table 1).

### Uncovering novel patterns in plant volatile responses

We applied the model to inducible volatile emissions in maize (*Zea mays*) in real time by PTR-ToF-MS following different stress treatments. First, we focused on the homoterpene 4,8-dimethylnona-1,3,7-triene (DMNT) as a highly inducible, ecologically relevant plant volatile (*71–73*). We tested whether the amount of mechanical leaf damage influences DMNT induction. Earlier work showed that damage intensity strongly correlates with terpene emissions (*12*). Indeed, higher damage increased total DMNT emissions (Fig. 2A), and our model faithfully captured this pattern through *Integral* (Fig 2B). No clear patterns were detected for the other parameters, apart from differences in *Duration*, which was marginally longer for intermediate amounts of wounding (Fig. 2D).

**Figure 2.**
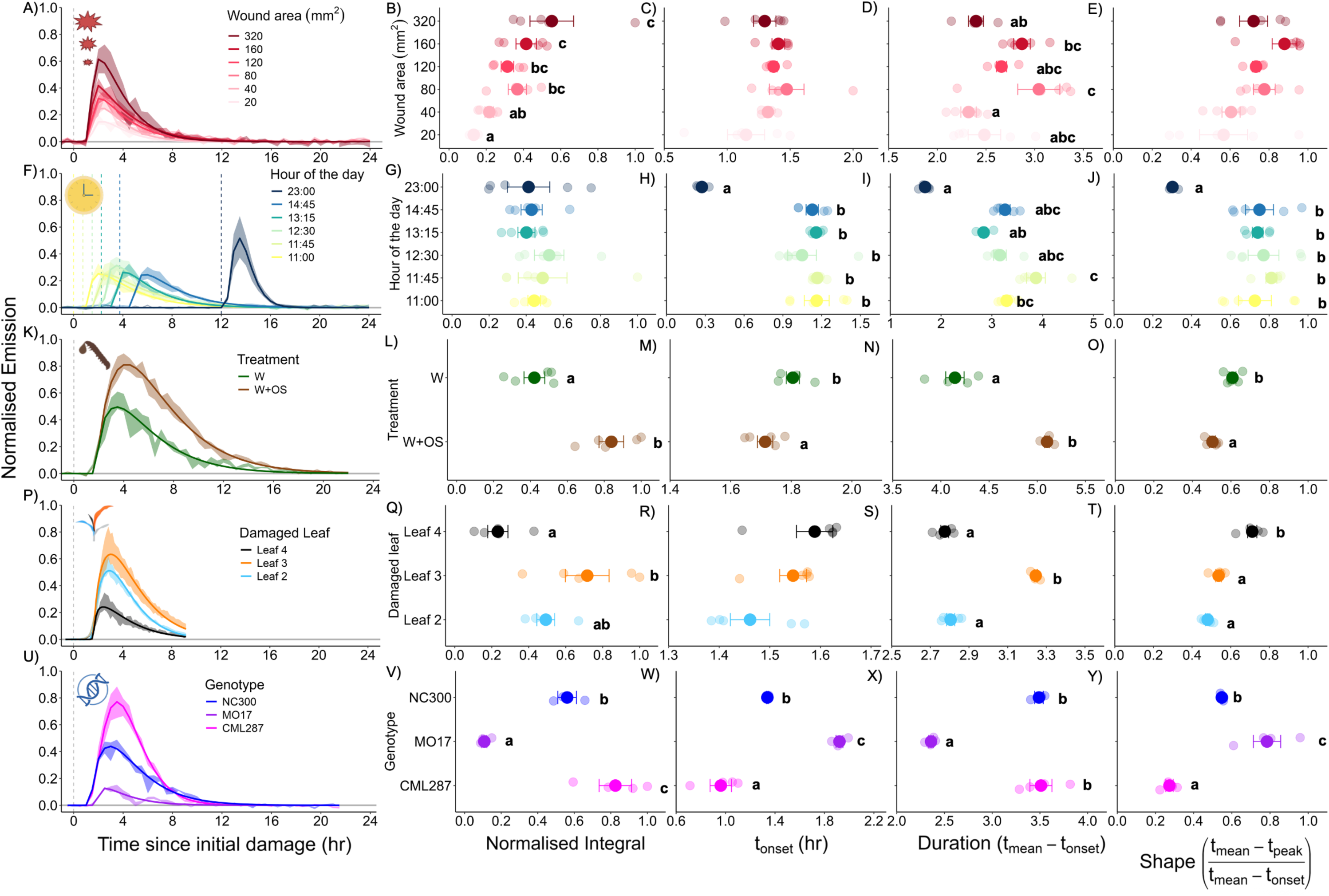
Application of the model to compare volatile emission dynamics in response to a range of stimuli. Each horizontal row of panels represents a unique experiment. A-E) responses to variable wounding intensities, F-J) responses to the same intensity of damage at different times of day under constant light (Note: *t_onset_* is standardised based on time of damage), K-O) responses compared between wounded plants and plants treated with wounding and *Spodoptera exigua* oral secretions (OS), P-T) responses to damage in leaves of different developmental stages (Data from Waterman et al., 2025), U-Y) wounding responses compared between genotypes with highly variable volatile emission capacity. For the first column of panels, curves depict emission data, where the solid line represents mean of fitted emission across biological replicates and the translucent ribbon represents the baseline-subtracted raw emission data ± SE. For the remaining columns, solid points represent mean values across biological replicates (translucent points). Error bars represent SE. Within13 each panel, different letters indicate significant differences between groups as determined by multiple comparisons tests following significant (*p* < 0.05) one-way tests *n* = 3-5.

Next, we tested the influence of the circadian clock on wound-induced DMNT by wounding plants at different times of day under continuous light. The circadian clock is known to play a role in regulating how plants respond to stress, including transcriptional regulation of defence genes (*74*, *75*). Time of day had no impact on the overall amount of DMNT emission (Fig 2G). However, *t_onset_*, *Duration* and *Shape* all varied significantly; wounding in the evening resulted in a significantly more rapid, shorter DMNT burst – a novel pattern that has not been reported before (Fig. 2F-J).

We also tested the influence of insect oral secretions (OS), which contain both elicitors and effectors which can increase and decrease responses compared to wounding alone, respectively (*76*). We confirmed that OS enhance the emission *Integral* of DMNT (Fig 2K-L). Interestingly, OS treatment also led to a more rapid and prolonged response, resulting in a significantly different curve shape (Fig 2M-O).

Leaf size and developmental stage can influence total volatile emissions (*57*), but whether these also influence response dynamics is unknown. We thus reanalysed the dataset from our earlier work with our model. We found that leaf 3 (largest size and intermediate age) emits volatiles for the longest period (Fig 2S). Interestingly, leaf 3 did not produce more DMNT than leaf 2 (older leaf) but produced it for a longer period of time (Fig 2Q and S). Both *t_onset_* and *Shape* followed a clear developmental gradient, whereby older leaves exhibited slower and more symmetrical emission dynamics compared to younger leaves (Fig 2R and 2T).

Different genotypes vary strongly in the amount and type of volatiles they emit (*77*), but whether response curves are also different is unknown. We measured response curves in three maize inbred lines that differ in their capacity to produce DMNT (Fig 2U). As expected, *Integral* varied for all genotypes (Fig 2V). Interestingly, we found that higher emitting genotypes also had substantially earlier *t_onset_* (Fig 2W), however there appears to be plasticity in the relationship between early onset and higher emissions as they are not always correlated (Fig 2A-C). Additionally, the two highest emitting genotypes, CML287 and NC300, had similar *Duration* (Fig 2X) and CML287 had the most symmetrical dynamics. Thus, different genotypes show starkly different response patterns, some of which (in this case *Duration*) appear conditionally independent of total volatile quantity. Taken together, these experiments illustrate the power of our approach to uncover novel, genetically determined response patterns to environmental stimuli.

### Unravelling differences between biochemically distinct compound classes

To explore how different volatiles respond to the same stimuli, we characterised volatiles from different biosynthetic pathways, including terpenes, indole and green leaf volatiles (*1*). In response to wounding, the different volatiles exhibited significantly different response curves, as described before, which we demonstrate quantitatively through *t_onset_*, *Duration* and *Shape* (Fig. 3A-E; Fig S1). Interestingly, the differences became more pronounced with the application of OS (Fig. 3F-J). This pattern was evident across all parameters but was strongest for *Duration*, whereby in the presence of OS each compound was produced for a unique amount of time, which was not the case in the absence of OS (Fig 3E and 3J). This illustrates how OS components can differentially modulate volatile pathways compared to mechanical wounding across volatile groups.

**Figure 3.**
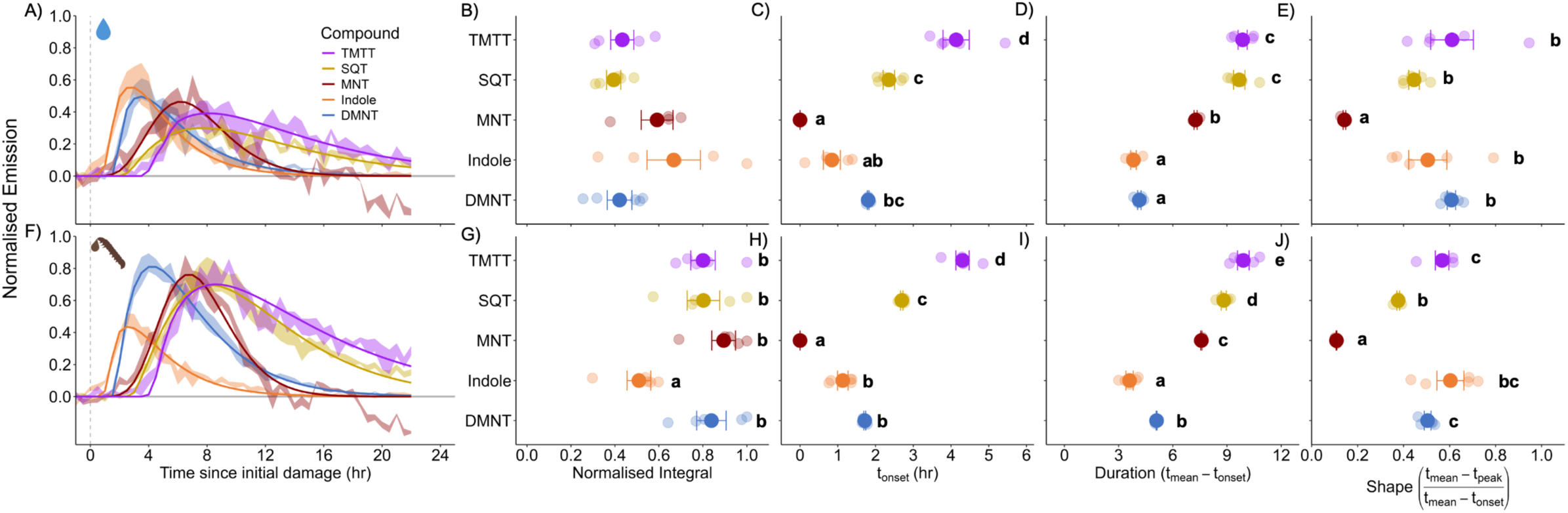
Impacts of herbivore-specific stimuli on volatile emission dynamics. Emission and model parameters for wounded plants (A-E) and wounded plants treated with oral secretions (OS; F-J). For the first column of panels (A and F), curves depict emission data, where the solid line represents mean of fitted emission across biological replicates and the translucent ribbon represents the baseline-subtracted raw emission data ± SE. For the remaining columns, solid points represent mean across biological replicates (translucent points). Error bars represent SE. Within each panel, different letters indicate significant differences between groups as determined by multiple comparisons tests following significant (*p* < 0.05) one-way or Welch’s ANOVA. *n* = 4-5. Abbreviations: DMNT= 4,8-dimethylnona-1,3,7-triene, MNT = monoterpenes, SQT = sesquiterpenes, TMTT = 4,8,12-trimethyltrideca-1,3,7,11-tetraene.

**Supplemental Figure 1.**
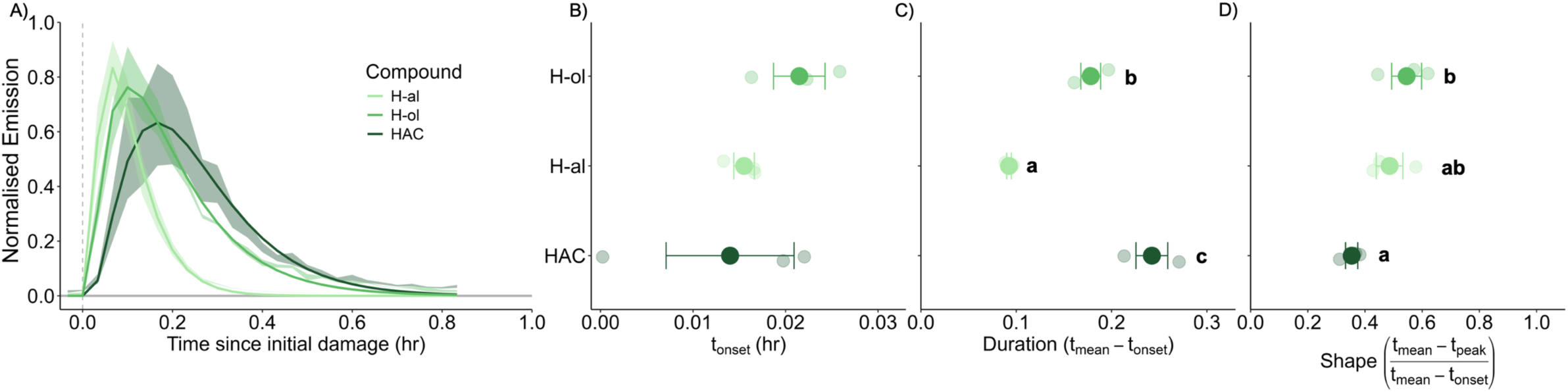
Dynamics of green leaf volatile (GLV) emissions. For A) curves depict emission data, where the solid line represents mean of fitted emission across biological replicates and the translucent ribbon represents the baseline-subtracted raw emission data ± SE. For the remaining columns, solid points represent mean across biological replicates (translucent points). Error bars represent SE. Within each panel, different letters indicate significant differences between groups as determined by multiple comparisons tests following significant (*p* < 0.05) one-way ANOVA. *n* = 3. Abbreviations: H-al = hexenal, H-ol = hexenol, HAC = hexenyl acetate.

To explore this phenomenon more deeply and test our model on a less temporally resolved dataset, we modelled the wound-responsive gene expression dynamics of *ZmCYP92C5*, *ZmIGL, ZmTPS2* and *ZmTPS10* (Fig S2), which are coding for the rate limiting enzymes of the biosynthesis pathways of DMNT/TMTT, indole, monoterpenes and sesquiterpenes, respectively (*57*). Trends in *Duration* of the expression of these genes matched emission of corresponding volatiles (Fig S2C and Fig 3E), confirming that volatile emission is, at least in part, regulated by biosynthetic limitations (Fig S2). There were clear differences in *t_onset_* detected (Fig S2B), however the lack of temporal resolution in measurements of gene expression dynamics makes clear interpretations of *t_onset_* difficult (Figs S3 and S4). Interestingly, although clear, differences in *Shape* did not match corresponding emission values (Fig S2D), suggesting unique parameterisation across levels of organisation. However this may be, at least partially, explained by low resolution.

**Supplemental Figure 2.**
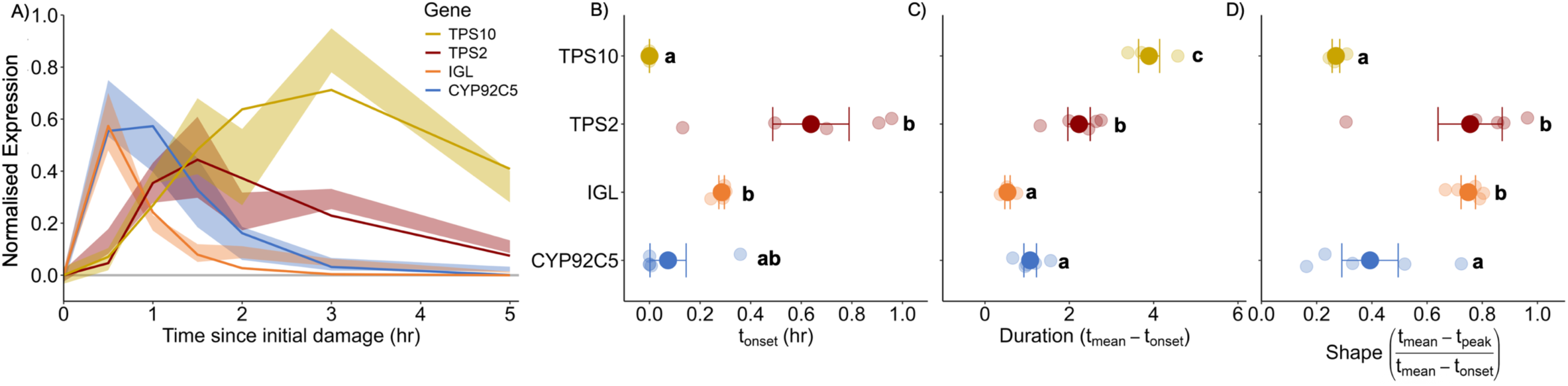
Volatile biosynthesis gene expression dynamics. For A), curves depict baseline-subtracted raw expression data, where the solid line represents mean of fitted expression across biological replicates and the translucent ribbon represents the baseline-subtracted raw expression data ± SE. For the remaining columns, solid points represent mean across biological replicates (translucent points). Error bars represent SE. Within each panel, different letters indicate significant differences between groups as determined by multiple comparisons tests following significant (*p* < 0.05) one-way ANOVA. *n* = 4-5. Abbreviations: CYP92C5 = dimethylnonatriene/trimethyltetradecatetraene synthase, IGL = indole-3-glycerol phosphate lyase, TPS2 = terpene synthase 2, TPS10 = terpene synthase 10 .

### Exploring complex damage and response patterns

To characterize more complex induction patterns, we quantified DMNT emission following multiple wounding events (Fig 4A). The current assumption from the literature is that subsequent wounding should lead to stronger defence responses beyond cumulative effects, as earlier wounding events will prime plants for subsequent responses. However, such effects have been hard to isolate for wounding events that follow each other closely in time due to overlapping response curves. Thus, we modelled the responses to three wounding events as the sum of three separate curves, all of which being effectively incomplete or not fully resolved in some way. By decomposing the overall emission into separate fitted curves, we were able to quantify the dynamic contribution of each individual wounding event, even when the peaks overlapped. This revealed a clear priming effect where the *integral* of the second and third responses were larger than that of the initial peak, demonstrating that prior damage enhanced subsequent emissions beyond additive effects (Fig 4B). At this particular time interval, multiple damage events did not significantly modify other fit parameters (Fig 4C-E). The functionality of this model on more complex damage patterns highlights its utility to explore response dynamics under complex stress regimes.

**Figure 4.**
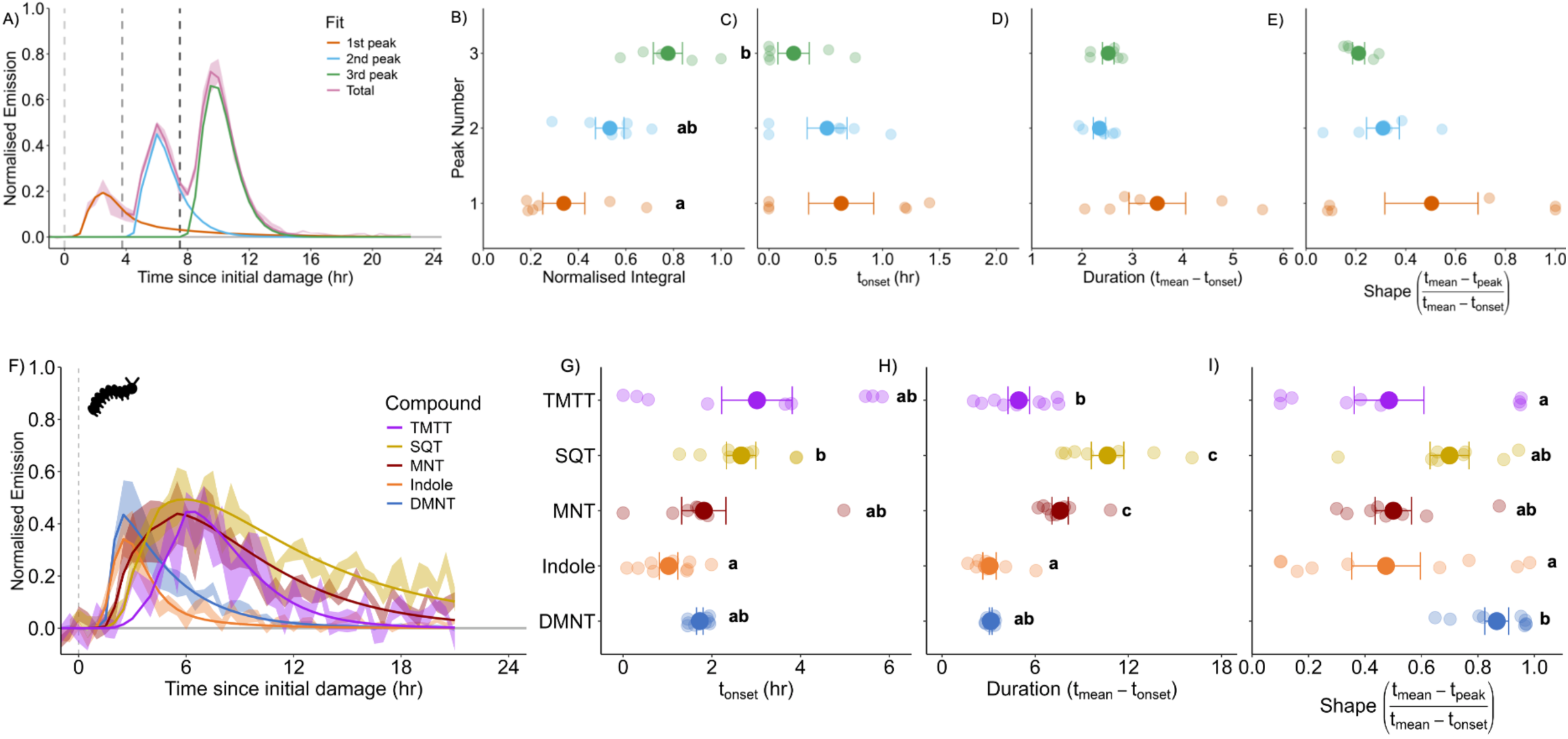
Fitting responses to complex stimulus patterns. A) Fitted DMNT emission from plants damaged three times (each 3.75 hr apart) for the total curve and each individual peak curves. B-E) Model parameters for each peak. F) Curves depict emission data and G-I depict model parameters from *Spodoptera exigua*-infested plants. For A and F, the translucent ribbons represent the baseline-subtracted raw emission data ± SE from the respective, colour-coded curve. For B-E and G-I, solid points represent mean across biological replicates (translucent points). Error bars represent SE. Within each panel, different letters indicate significant differences between groups as determined by multiple comparisons tests following significant one-way ANOVA. For A-E, *n* = 6 and for F-I, *n* = 8-9. Abbreviations: DMNT= 4,8-dimethylnona-1,3,7-triene, MNT = monoterpenes, SQT = sesquiterpenes, TMTT = 4,8,12-trimethyltrideca-1,3,7,11-tetraene.

Herbivores do not make single wounds when they feed, but rather take many bites over time. As such, we quantified emission responses to feeding by a single *Spodoptera exigua* larva for 30 min. Natural herbivory patterns generated responses that could be easily fit for all groups of volatiles (Fig 4F). *t_onset_* showed some similarities to wounding response patterns, namely that sesquiterpenes and TMTT take longer to begin emitting than other compounds (Fig 4G). Interestingly, *Duration* values more closely matched those from mechanical wounding than wounding + OS, with fewer unique durations forming (Fig 4H). *Shape* values showed stark differences compared to wounding treatments, suggesting that damage patterns may play an important role in determining response curves (Fig 4I). The ability to effectively fit parameters to responses induced by real herbivore feeding indicates that our model can be used to detect differences in response dynamics even when exact damage patterns and timing are not known.

### Quantifying incomplete curves

Understanding the dynamics of defence responses requires some level of continuous monitoring. However, how the temporal resolution of measurements impacts the capacity to understand dynamic patterns is not well known. We tested how measurement duration and sampling resolution affected quantification. Firstly, in order to simulate a shorter measurement window, we removed datapoints from the end of the curve (in time) and compared fit parameters to those from the complete dataset. For *Integral* and *Duration*, there was a consistent point around *t_mean_* where removal of data dramatically reduced the accuracy of model parameters (Fig S3B, S3D, S3G, S3I, S3L and S3N). *t_onset_* was more sensitive to data removal for indole and TMTT than it was for DMNT (Fig S3C, H and M). Similarly for *Shape* (which is *t_onset_*-dependent), indole and TMTT fits broke down relative to the complete dataset with less data removal than DMNT (Fig S3E, J and O). Interestingly, for DMNT and indole, even with removal of *ca.* 10-15 hr of curve the fitted parameters remained almost entirely stable, if not identical, suggesting a substantial degree of flexibility and potential to understand the dynamics of only partially resolved peaks.

Second, we tested the resolution tolerance of the model by periodically removing datapoints, simulating a lower resolution measurement (Fig S4A, F and K). With periodic removal of up to 85% of data points, the emission *Integral* was able well recovered for all compounds, suggesting that a coarse ‘outline’ of temporal response patterns can be sufficient to accurately estimate total emissions over time (Fig S4B, G and L). For indole, removal of 70% data in periodic intervals resulted in breakdown of *Integral* as the overall curve structure was nearly entirely lost (Fig S4B). Indole has the fastest emission dynamics, and as such, has the lowest effective resolution in comparison to DMNT and TMTT, which both have slower kinetics and thus more data points between *t_onset_* and *t_mean_*. Interestingly, other fit parameters (*t_onset_*, *Duration* and *Shape*) began to consistently breakdown at 70% removal, suggesting there is a limit to measurement frequency for certain dynamic features (Fig S4). Importantly, these results highlight that response kinetics are a necessary consideration to determine the use of truly ‘continuous’ measurements; they may not always be required or even provide additional, useful information.

**Supplemental Figure 3.**
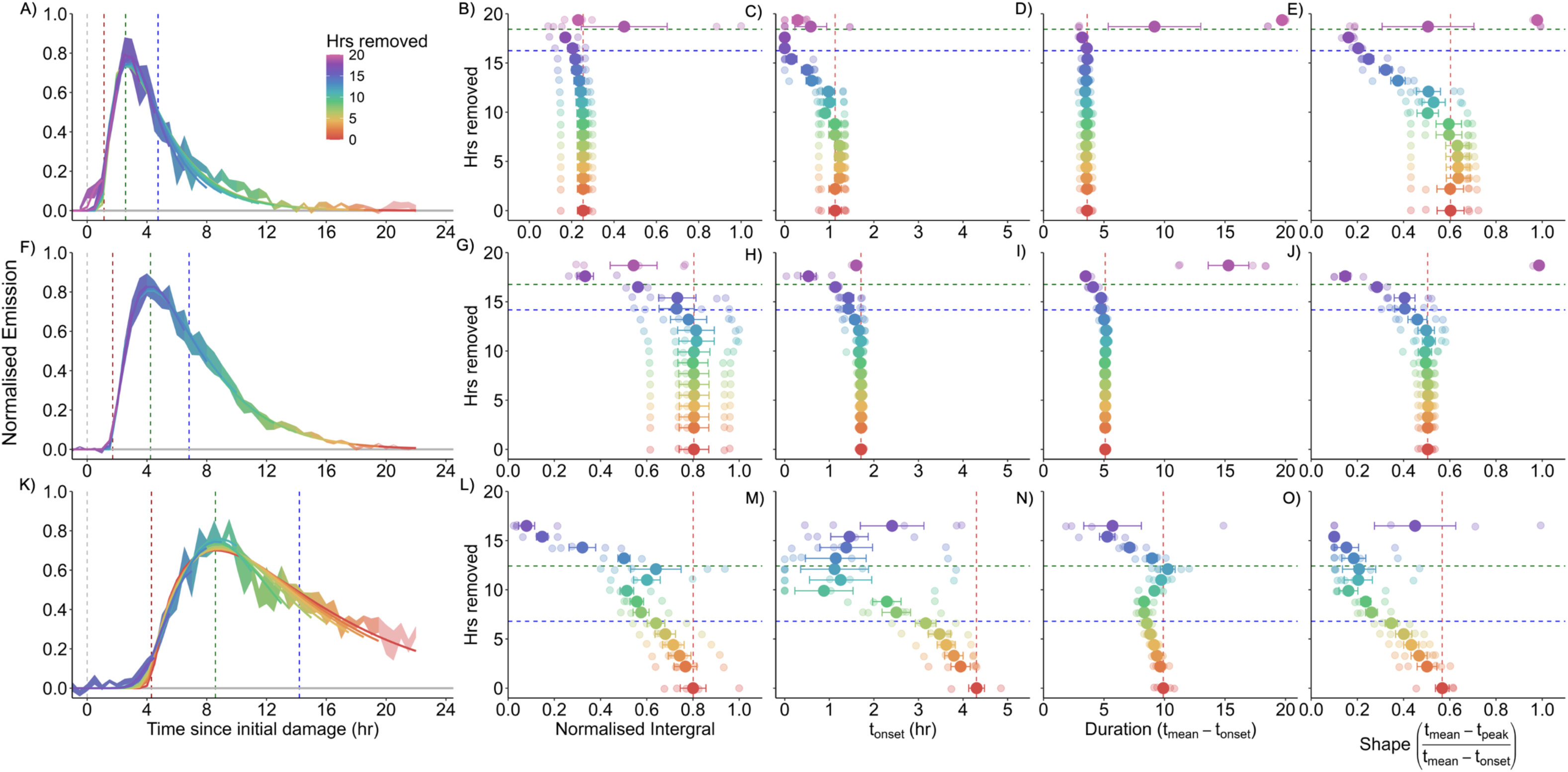
Fitting incomplete response curves. Emission and fit parameters for A-E) indole, F-J) DMNT and K-O) TMTT. For the first column of figures, curves depict emission data, where the each solid line represents mean of fitted emission across biological replicates and the translucent ribbon represents the raw baseline-subtracted emission data ± SE. For the remaining columns, solid points represent mean across biological replicates (translucent points). Error bars represent SE. Colour gradient indicates the number of hours removed from the back end of the curve. *n* = 5. Abbreviations: DMNT= 4,8-dimethylnona-1,3,7-triene, TMTT = 4,8,12-trimethyltrideca-1,3,7,11-tetraene.

**Supplemental Figure 4.**
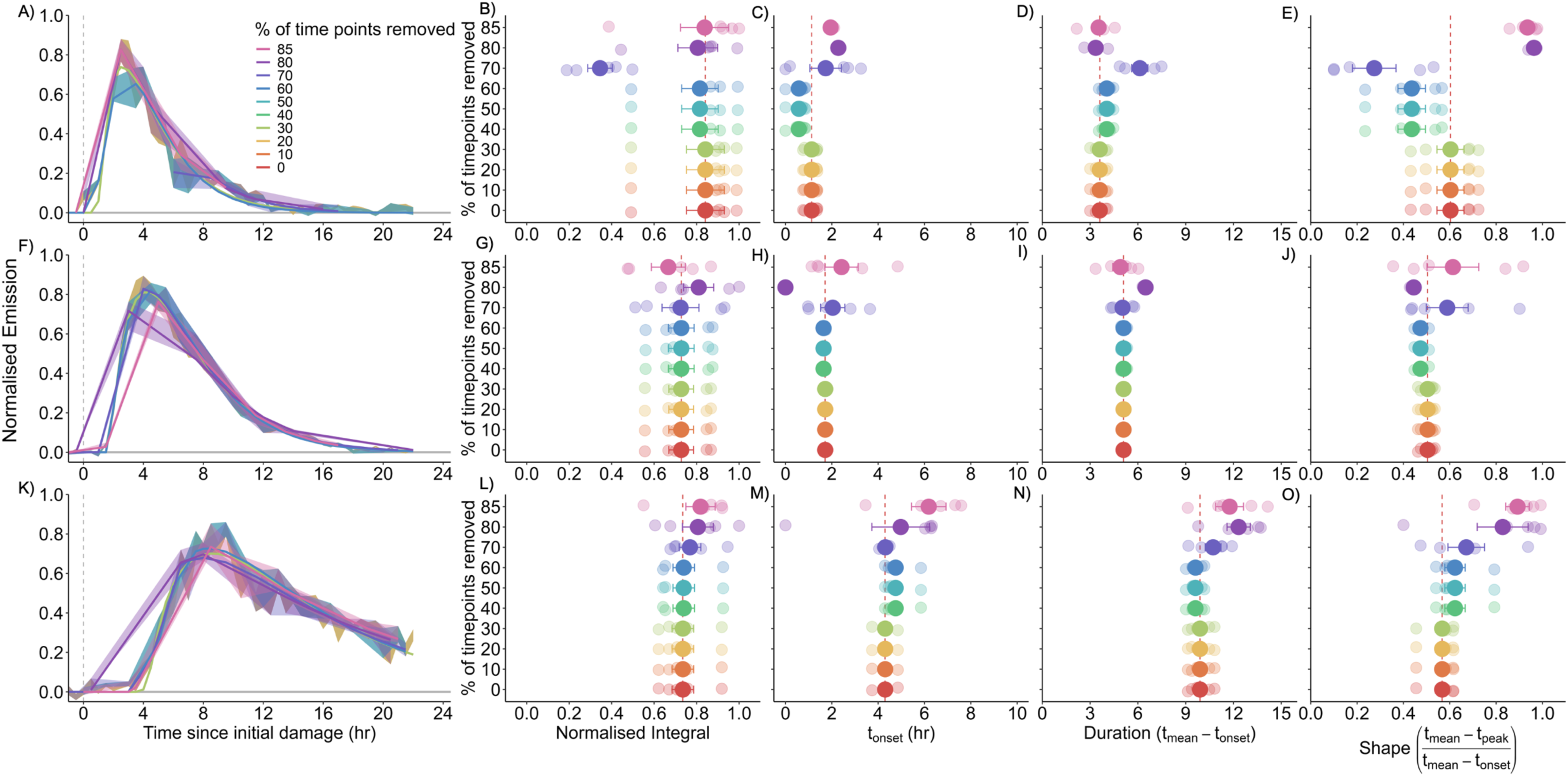
Sampling resolution impacts fits. Data were removed systematically in periodic intervals across the entire curve. Emission and fit parameters for A-E) indole, F-J) DMNT and K-O) TMTT. For the first row of data, curves depict emission data, where the solid line represents mean of fitted emission across biological replicates and the translucent ribbon represents the baseline-subtracted raw emission data ± SE. For the remaining columns, solid points represent mean across biological replicates (translucent points). Error bars represent SE. *n* = 5. Abbreviations: DMNT= 4,8-dimethylnona-1,3,7-triene, TMTT = 4,8,12-trimethyltrideca-1,3,7,11-tetraene.

### Applying the framework across broader systems and response types

To demonstrate the wide application of our framework and the breadth of consistent curve structure in inducible responses across the tree of life, we applied it to response dynamics following different stimuli across diverse plant and animal systems, measurement techniques and nearly seven orders of magnitude of time (*51–56*). These systems all conformed well to the overall gamma structure enabling meaningful parameter extraction (Fig S5A-E). Our approach applies a universal language of identical parameters across systems, enabling comparison and quantitatively elucidation of differences in response dynamics between groups.

**Supplemental Figure 5.**
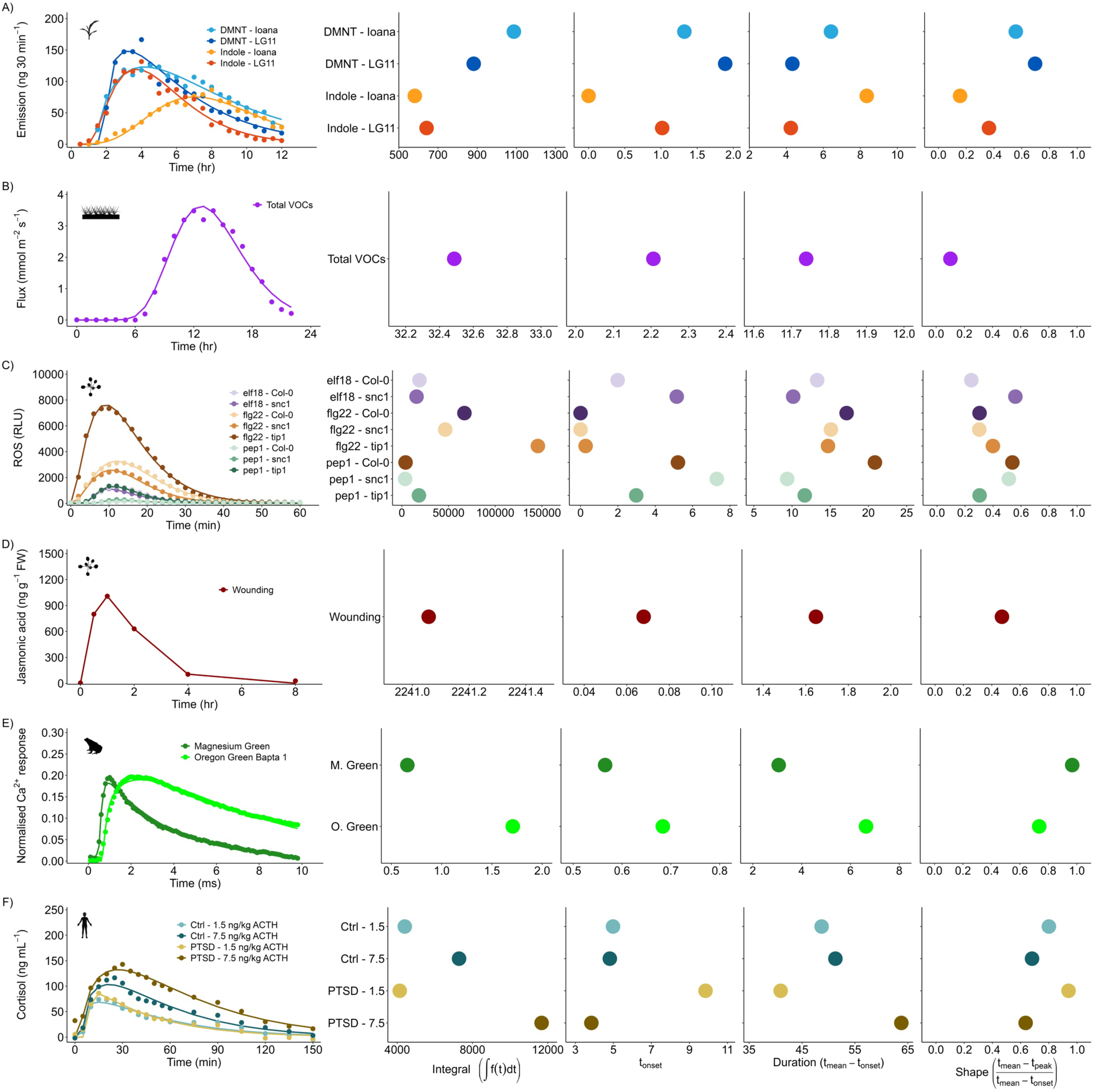
The framework can be used to fit and parameterise the temporal dynamics of responses across organisms and response types. A) DMNT and indole emission in two maize genotypes (Ioana and LG11) measured by GC-MS from Turlings et al., 1998. B) Open-headspace VOC emission fluxes in a winter wheat field measured by PTR-Qi-ToF-MS from Loubet et al., 2022. C) Reactive oxygen species (ROS) burst in response to two bacterial peptide elicitors (elf18 and flg22) and one plant-secreted elicitor (pep1) in three *Arabidopsis thaliana* genotypes: wildtype (Col-0), an autoimmune mutant (snc1) and tip growth defective 1 (tip1) from Cheng et al., 2024. D) Jasmonic acid production in response to mechanical wounding in *A. thaliana* from Park et al., 2002. E) Calcium transients measured by two different fluorescent dyes (Magnesium Green and Oregon Green Bapta 1) in marsh frog (*Rana ridibunda*) nerve terminals in response to an electrical stimulus from Samigullin et al 2015. F) Baseline-subtracted cortisol response in human subjects with a PTSD diagnosis (PTSD) or no psychiatric disorder (Ctrl) in response to blood infusion with adrenocorticotropic hormone (ACTH) at differing concentrations from Radant et al 2009. In the left-most column, coloured points represent raw data extracted from respective studies and lines represent individual fitted response values. Points in parameter figures represent modelled values for respective parameters.

## Discussion

Dynamic responses to environmental stimuli are a universal property of life. Yet, unifying approaches to characterize such responses are lacking. Across biology, we have major gaps in our understanding of patterns and differences in dynamic biological responses. Induced volatile emissions are an illustrative example in this context. In a series of experiments, we find that our model reveals novel differences in volatile induction along circadian, developmental and genetic axes, and in response to wounding and insect-specific molecular patterns. Additionally, we uncouple convoluted response patterns to identify rapid priming events that can inform how complex stress regimes might change defences trajectories compared to simple ones. Among the most noteworthy discoveries is that, independent of light, the time of day of wounding has a strong impact on the onset, duration and shape of the volatile induction curve but not the total response intensity. This provides a window into the tight regulation of the speed and duration of induced volatile emissions by the circadian clock (*10*, *78*), providing new phenotypes that link clock regulation to environmental interactions. Equally noteworthy is the finding that insect oral secretions (OS) enhance differences in response curves between different volatile classes, thus resulting in unique temporal volatile fingerprints. This newly discovered pattern may explain OS-specific responses that affect interactions with herbivores and herbivore natural enemies beyond differences in overall volatile quantities or timing (*79*). Finally, we uncover that, contrary to current expectations, the quantity and duration of induced volatile emissions can be conditionally independent across plant genotypes. This finding paves the way to identify regulators of volatile induction duration through genetic approaches, potentially leading to the identification of new “late signalling components” involved in sustaining defence responses. Further, these novel dynamic features might ultimately be leveraged to non-destructively distinguish and identify stimuli, for example in an agricultural context, where many individual induced responses may be shared between multiple stressors, and total response is thus insufficiently informative (*6*, *47*, *58*).

Recent advances in continuous, real-time monitoring have greatly improved our ability to measure inducible response dynamics (*5*, *7*, *10*, *58*), yet quantitative analyses of such data remain limited. Our modelling approach addresses this gap by providing a standardised framework for quantifying response dynamics in a way that enables biologically and ecologically meaningful comparisons across systems. In this study, we use volatile emissions and gene expression from plants as representative measures of functionally and biochemically diverse inducible responses. Additionally, we use our framework on six additional examples from published studies. And while we do not draw any biological conclusions from these studies, we demonstrate the utility of our framework across biological systems, from plants to humans (*51–56*). Thereby, the framework opens opportunities for ambitious endeavours, such as tree-of-life scale meta-analyses, to uncover shared and divergent principles of organismal responses to environmental stimuli and to reveal new fundamental biological phenomena. Going forward, coupling model parameters with machine-learning algorithms or constrained optimisation solvers could enable the reconstruction of stimulus patterns from observed dynamics, or the prediction of dynamic responses from environmental inputs. We propose this framework as a universal standard for intuitive, interpretable and biologically grounded quantification of inducible responses, to advance understanding of fundamental processes and to guide innovation across medicine, agriculture and beyond.

To ensure that the model can be used widely, we generated a set of freely accessible resources that can be implemented in Python, R and Excel formats (Appendix I), and applied broadly to time-resolved response data to yield all relevant readouts for further statistical analyses. Additionally, our entire analytical pipeline, including additional model details can be found here: https://github.com/watermaj/Quant-dynamics.

## Materials and Methods

### Plants and insects

Plant growth conditions were identical to those used in Waterman et al. (2025). In brief, V2-stage (two developed leaves, one expanding leaf and one emerging leaf) maize (*Zea* mays) seedlings were used throughout the study. At this stage maize plants are particularly susceptible to agricultural pests such as *Spodoptera* spp. (*80*). Plants were grown in commercial potting soil (Selmaterra, BiglerSamen, Switzerland) in 180 mL pots under greenhouse conditions and supplemented with artificial lights (ca. 300 µmol m−2 s−1). The greenhouse was kept at 22 ± 2°C, 40%–60% relative humidity, with a 14 h: 10 h, light: dark cycle.

*Spodoptera exigua* larvae (Frontier Agricultural Sciences, USA) were reared from eggs on an artificial diet (*81*). At least 24 hr prior to experimental treatments larvae were fed B73 maize leaves. Oral secretions (OS) were collected by probing the mouths of larvae with a pipette tip and stored at -20°C until use.

### Experimental Treatments

All wounds were inflicted using haemostat forceps, an established method of simulating herbivory (*57*, *76*). The maize inbred line, B73, was used unless otherwise stated. The standard damage intensity was 120 mm^2^ and time of initial damage was 11:00 unless otherwise stated. To understand how different damage regimes impact induced response dynamics we conducted several experiments spanning a range of treatments:

#### Variable damage intensity

We damaged plants at 20, 40, 80, 120, 160 and 320 mm^2^ on the third-oldest leaf (leaf 3). Damage patterns always encompassed the central segment of each leaf, and any damage above 40 mm^2^ was evenly distributed across the base, middle and tip of leaf 3 (*57*).

#### Time of day

We damaged plants at 11:00, 11:45, 12:30, 13:15, 14:45, 15:45 and 23:00.

#### Oral secretions (OS)

Damaged plants were either treated with milliQ water or 50% OS from late-instar *Spodoptera exigua* larvae.

#### Leaf developmental stage

At the V2 stage maize has four leaves: two are fully developed (leaf 1 and leaf 2), one is actively expanding (leaf 3) and one is emerging (leaf 4). We used the raw emission data from Waterman et al. (2025) to measure emission dynamics in each leaf. The damage treatments in the previous study were 60mm^2^ total, and damage was inflicted in three bouts of 20 mm^2^ damage in *ca.* 30 min intervals

#### Genotype

To explore the potential variation in response dynamics across maize genotypes we damaged three additional maize inbred lines, with three distinct volatile emission capacities: MO17 (low emission), NC300 (intermediate-high emission) and CML287 (high emission).

#### Real herbivory

3^rd^ instar *Spodoptera exigua* were placed on leaf 3 and were left to feed for 30 minutes.

### Volatile sampling

Volatiles were collected as in Waterman et al. (2025). Briefly, entire seedlings were placed in transparent glass chambers (Ø×H 12 × 45 cm) that were sealed other than a clean airflow inlet and an outlet. Clean air was supplied at a flow-rate of 0.8 L min^−1^. Volatiles were measured with a high-throughput platform comprising of a proton transfer reaction time-of-flight mass spectrometer (PTR-ToF-MS; Tofwerk, Switzerland) and a custom-made automated headspace sampling system. The outlet of the chamber was accessible to the autosampler/PTR-ToF-MS system, which drew air at 0.1 L min^−1^. Between samples, a zero-gas measurement was performed for 3 s to flush the system. At each time point (as indicated by the x-axis of the respective figure), volatiles were continuously measured for 25–30 s and averaged to a single mean per sample. Complete mass spectra (0–500 m/z) were recorded in positive mode at *ca*. 10 Hz. The PTR was operated at 100°C and an E/N of approximately 120 Td. The volatile data extraction and processing were conducted using Tofware software package v3.2.2 (Tofwerk, Switzerland). Protonated compounds were identified based on their molecular weight + 1. During volatile collection, LED lights (DYNA, heliospectra, Sweden) were placed *ca*. 80 cm above the glass chambers and provided light at 300 μmol m−2 s−1. Identical light:dark cycle timing as in the greenhouse for plant growth was used unless otherwise stated.

### Gene expression

Leaf 3 tissue was harvested 0.5, 1, 1.5, 2, 3 and 5 hr after damage (see above for damage treatment) and flash frozen on liquid nitrogen. Total RNA extraction and purification, genomic DNA removal, cDNA synthesis and quantification of gene expression were conducted identically to Waterman et al. 2024. Quantitative reverse transcription polymerase chain reaction (qRT-PCR) was performed using ORA SEE qPCR Mix (highQu GmbH, Germany) on an Applied Biosystems QuantStudio 5 Real-Time PCR system. The normalised expression (NE) values were calculated as in Waterman et al. (2024) using ubiquitin (UBI1) as the reference gene. Gene identifiers and primer sequences are listed in Table S1.

### Modelling emission dynamics

For all experiments, volatile emissions were normalised by the biomass of leaf 3 (damaged leaf). This is because we have previously shown that the size of the damaged leaf is limiting for volatile emissions (*57*).

#### Choosing the right distribution

To model the waiting time probability distribution, we used a Gamma distribution:

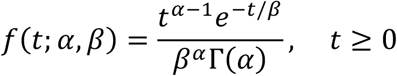

Where:

• *α* is the number of steps,

• *β* is the rate parameter (for each step),

• Γ(*α*) is the Gamma function evaluated at *α*

This distribution arises from the sum of several exponential waiting times, making it suited to processes where multiple sequential steps precede a single observable outcome (*82*). The Gamma distribution is closed under addition, meaning that the sum of sequential Gamma is itself a Gamma distribution (*83*). This allows modelling of both early and late phase responses, even if they are both part of the same chain of events. In the formal definition, *α* represents the effective number of steps and *β* an effective average delay, though we do not assume that these biological processes are truly memoryless and independent as required by the strict derivation. Instead, we treat the Gamma distribution as a phenomenological approximation of the process, which fits the shapes of the responses we (and others) observe well (*1*, *2*, *28*, *57–65*). With this in mind, we make a number of meaningful reparameterisations to the standard Gamma distribution, as follows.

We begin with the standard Gamma distribution and introduce a shifted time variable *t_adj_* to account for a delay in the onset of the curve:

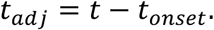

The Gamma probability density function is then given by:

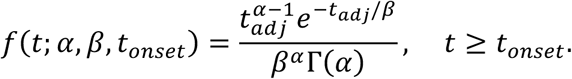

Our next step is to relate the variables to observable quantities, and so we want to express *α* and *β* in terms of measurable features of the response curve. For the Gamma distribution:

_•_ *Mode* = (*α* − 1)*β* = *t_peak_* − *t_onset_*

representing the time at which the response reaches its peak.

_•_ *Mean* = *αβ* = *t_mean_* − *t_onset_*

representing the average time after onset that the response occurs.

In order to solve for *α* and *β* for these observable features we: Subtract the first from the second:

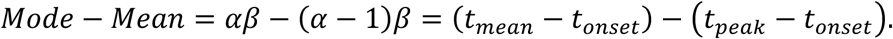

Simplify:

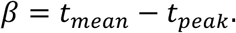

Substitute into *αβ* = *t_mean_* − *t_onset_*:

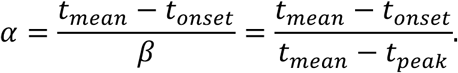

The Gamma probability density function is normalised to integrate to 1.

However, in experimental data the absolute amplitude (the response intensity, *R_peak_*) is an observable quantity and so we define the response model:

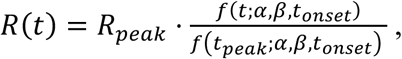

where *R_peak_* is the observed response at *t_peak_*.

This ensures that the curve shape follows the Gamma form, and that the maximum of *R*(*t*) equals *R_peak_*.

The next step is to solve for *f*-*t_peak_*; *α*, *β*, *t_onset_*.

Substitute *t* = *t_peak_* into *f*(*t*; *α*, *β*, *t_onset_*):

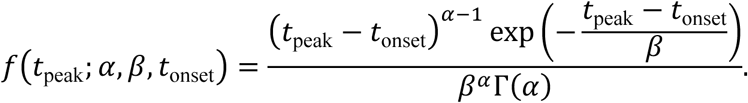

Replace *t_peak_* − *t_onset_* with (*α* − 1)*β* using the mode definition:

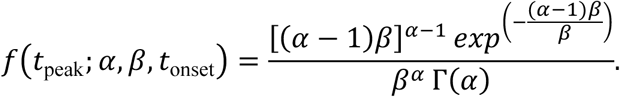

Simplify the exponent:

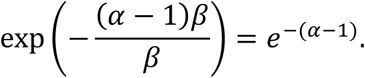

Final expression:

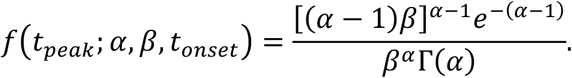

We can therefore define:

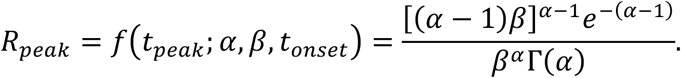

Substitute *f*(*t*)and *f*(*t_peak_*). into the ratio:

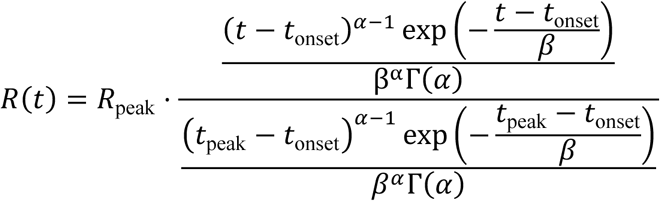

Cancel the common terms *β^α^*. Γ(*α*):

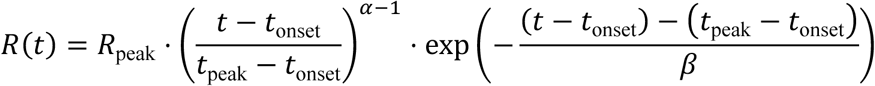

Simplify the exponent:

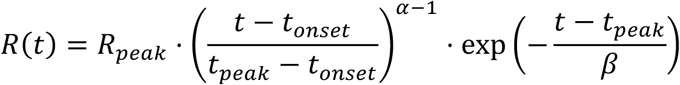

Replace α and β with expressions in terms of -*t_onset_*, *t_peak_*, *t_mean_*. to give the final model:

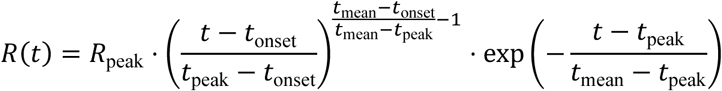

Where:

• *R_peak_* is the maximum measured response,

• *t_onset_* is the onset delay,

• *t_peak_* is the time of peak response,

• *t_mean_* is the mean response time.

#### Background subtraction

Since low levels of plant volatiles are constantly emitted even without wounding or other measurable stimuli, a background curve was subtracted so that only induced emissions were modelled (*44*). This background was estimated from undamaged plants and either directly subtracted or scaled to the pre-damage baseline to correct for instrumental drift or individual plant batch variation.

#### Fitting

Each individual curve was fit using a two-stage optimisation approach. In order to avoid local minima, we first used a differential evolution optimisation followed by a local refinement step using SLSQP (*84*). Since the emission data were individually relatively noisy, we reconstructed the dominant temporal trend using singular value decomposition (SVD) and incorporated this as a weak prior to guide the fitting algorithm toward biologically realistic solutions (*85*, *86*). A number of constraints and bounds were enforced in order to guide the fitting of the data. Temporal ordering was imposed such that *t_onset_* < *t_peak_* < *t_mean_*. To prevent degenerate solutions, minimum separations were required, with *t_peak_* − *t_onset_* > 0.1 and *t_mean_* − *t_peak_* > 0.1. Bounds were dynamically set for each curve; for instance, *t_peak_* was initialised at the time of maximum observed response and restricted to within ±20% of this value, while *R_peak_* was constrained between 80% and 120% of the observed maximum. *t_onset_* and *t_mean_* times were loosely defined based on empirically informed ranges so fitted values were virtually unconstrained. All fitting, calculations and data manipulation were done using SciPy (*68*), NumPy (*87*) and Pandas (*88*) in Python 3.12.4 (*89*).

To judge how well the model described the measured data, we calculated the coefficient of determination (R²) and the root mean squared error (RMSE) for every fitted curve. RMSE was normalised to the fitted peak height (*R_peak_*), so that error is expressed as a fraction of the response amplitude and is comparable between responses measured on different scales. These statistics were calculated at two levels. First, each fitted curve was compared with its own observed data (’individual fits’ in Supplemental Table 1). Individual replicates carry substantial biological and measurement noise, however, so these values conflate two sources of error: failure of the model to describe the response, and scatter that no smooth curve could follow. To separate the two, we repeated the analysis on averaged curves, comparing the mean observed emission with the mean fitted curve within each matched treatment group (’mean response curve’ in Supplemental Table 1). Averaging suppresses replicate-level noise while leaving the underlying response shape intact, so agreement here reflects how well the model captures the response itself.

#### Model testing

To explore the robustness of the model and fitting methods to the effects of resolution and measurement time, a set of data were removed in order to simulate a lower quality measurement. We chose the dataset of damage plants treated with insect OS, as a representative and ecologically relevant subset. In a first test, data from the end of the measurement were removed in a stepwise manner to simulate a shorter (incomplete) experimental measurement period. Secondly, data were systematically down-sampled to simulate a lower resolution measurement. After each step of the truncation or down sampling the data was fit and the fit parameters compared.

### Additional statistical analyses

All further statistical analyses were conducted in R version 4.4.2 (*90*). Differences in model outputs between groups were determined using one-way ANOVA. Where necessary, data were transformed to meet ANOVA assumptions. Where necessary, to obtain heteroscedasticity-consistent standard errors, White-adjusted ANOVAs were used (*91*).Where data did not meet normality assumptions, Kruskal-Wallis tests were used. Where data did not meet the homogeneity of variance and normality assumptions, even following transformations, differences were determined using Welch’s ANOVA. Statistical test summaries are included in Supplemental Tables 2-5. As emissions were expressed as ion counts in arbitrary units, observed and fitted emission values were normalised to the maximum observed response within each experiment for visual clarity, yielding positive values ≤ 1. Similarly, calculated response integrals were normalised to the maximum integral within each experiment. All normalisation was performed after model fitting and thus did not affect model fitting or parameter estimation.

### Data extraction from published studies

Data were extracted from published studies using WepPlotDigitiser (*92*).

## Acknowledgements

We would like to thank Sarah Holder for assistance with plant growth and Dr. Tristan Cofer for experimental assistance. This work was supported by the Swiss National Science Foundation (Grant Nr. 201651), the State Secretariat for Education, Research and Innovation (CANWAS), the University of Bern and Trinity College Dublin.

## Author contributions

Conceptualization: JMW, GM, LAC, ME

Methodology: JMW, GM, LAC, ME

Investigation: JMW, GM, LAC, SH, ME

Visualization: JMW, GM

Funding acquisition: JMW, ME

Project administration: JMW, ME

Writing (all drafts): JMW, GM, LAC, ME

## Data and Materials Availability

The Python scripts used to model response dynamics and the datasets are available at the GitHub repository: https://github.com/watermaj/Quant-dynamics. Additionally, we provide examples of how to adopt the presented approach in Python, R and Excel formats (see Appendix I).

## Supplemental Tables

**Supplemental Table 1.**
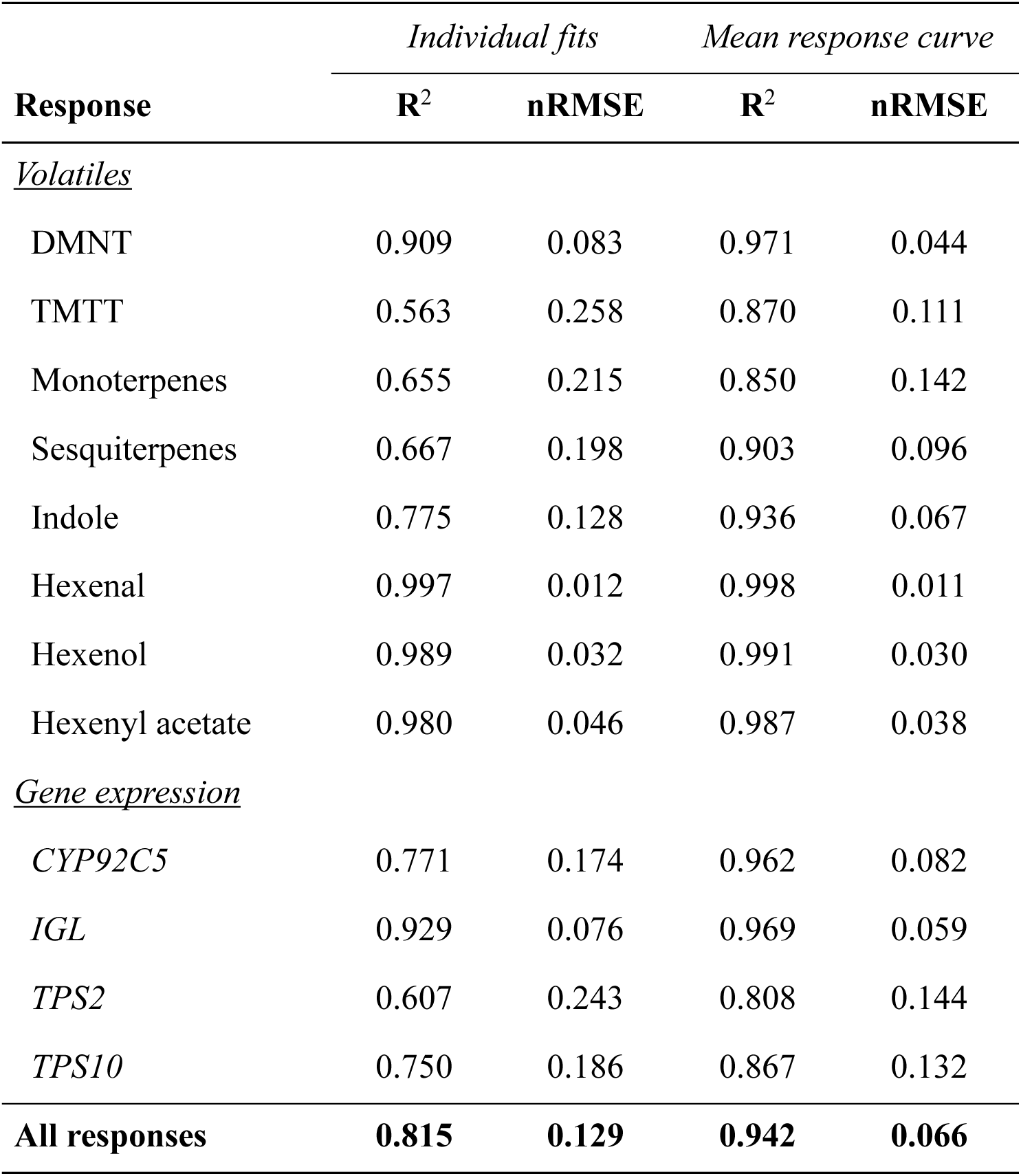
Goodness of fit of the model across all fitted responses. R^2^ and normalised root mean squared error (nRMSE) are reported for individual fitted curves (averaged within each response) and for the mean fitted curve compared with the mean observed curve within matched treatment groups. nRMSE is normalised to the fitted peak height, Rpeak. Sensitivity-analysis datasets (resolution-adjusted and time-truncated) are excluded.

**Supplemental Table 2.** Summary of statistical analyses presented in Figure 2 (main text). Bold values: *p* < 0.05. Underlined values: *p* <0.1. a = analysed using one-way ANOVA, b = analysed using Kruskal-Wallis test. * = analysed on log-transformed data, † = analysed using white-adjusted ANOVA.

| Response | *F* | *p-value* | *df* |
| --- | --- | --- | --- |
| <u>*Damage Intensity*</u> |  |  |  |
| *Integral*<sup>a*†</sup> | **20.70** | **< 0.001** | **5** |
| *t<sub>onset</sub>*<sup>a</sup> | 1.39 | 0.262 | 5 |
| *Duration*<sup>a*</sup> | **4.63** | **0.004** | **5** |
| *Shape*<sup>a</sup> | <u>2.60</u> | <u>0.051</u> | <u>5</u> |
| <u>*Circadian clock*</u> |  |  |  |
| *Integral*<sup>a*</sup> | 0.461 | 0.801 | 5 |
| *t<sub>onset</sub>*<sup>a*</sup> | **92.55** | **< 0.001** | **5** |
| *Duration*<sup>b</sup> | **24.46** | **< 0.001** | **5** |
| *Shape*<sup>a</sup> | **10.57** | **< 0.001** | **5** |
| <u>*Oral secretions*</u> |  |  |  |
| *Integral*<sup>a</sup> | **22.84** | **0.001** | **1** |
| *t<sub>onset</sub>*<sup>a</sup> | **7.49** | **0.026** | **1** |
| *Duration*<sup>a</sup> | **94.53** | **< 0.001** | **1** |
| *Shape*<sup>a</sup> | **19.53** | **0.002** | **1** |
| <u>*Leaf number*</u> |  |  |  |
| *Integral*<sup>a</sup> | **9.11** | **0.004** | **2** |
| *t<sub>onset</sub>*<sup>a</sup> | <u>3.61</u> | <u>0.060</u> | <u>2</u> |
| *Duration*<sup>a</sup> | **226.25** | **< 0.001** | **2** |
| *Shape*<sup>a</sup> | **46.59** | **< 0.001** | **2** |
| <u>*Genotype*</u> |  |  |  |
| *Integral*<sup>a†</sup> | **46.93** | **< 0.001** | **2** |
| *t<sub>onset</sub>*<sup>a†</sup> | **165.60** | **< 0.001** | **2** |
| *Duration*<sup>a</sup> | **74.39** | **< 0.001** | **2** |
| *Shape*<sup>a†</sup> | **76.79** | **< 0.001** | **2** |

**Supplemental Table 3.** Summary of statistical analyses presented in Figure 3 (main text). Bold values: *p* < 0.05. a = analysed using one-way ANOVA, b = analysed using Welch’s ANOVA, * = analysed on log-transformed data, † = analysed using white-adjusted ANOVA.

| Response | *F* | *p-value* | *df* |
| --- | --- | --- | --- |
| <u>*Without oral secretions*</u> |  |  |  |
| *Integral*<sup>†a</sup> | 1.94 | 0.145 | 4 |
| *t<sub>onset</sub>*<sup>b</sup> | **1465.30** | **< 0.001** | **4** |
| *Duration*<sup>†a</sup> | **231.49** | **< 0.001** | **4** |
| *Shape*<sup>*a</sup> | **29.71** | **< 0.001** | **4** |
| <u>*With oral secretions*</u> |  |  |  |
| *Integral*<sup>a</sup> | **5.99** | **0.002** | **4** |
| *t<sub>onset</sub>*<sup>b</sup> | **2328.30** | **< 0.001** | **4** |
| *Duration*<sup>a</sup> | **206.99** | **< 0.001** | **4** |
| *Shape*<sup>b</sup> | **561.00** | **< 0.001** | **4** |

**Supplemental Table 4.** Summary of statistical analyses presented in Figure 4 (main text). Bold values: *p* < 0.05 and underlined values: *p* < 0.1. a = analysed using one-way ANOVA, b = analysed using Welch’s ANOVA, * = analysed on log-transformed data, † = analysed using white-adjusted ANOVAs.

| Response | *F* | *p-value* | *df* |
| --- | --- | --- | --- |
| <u>*Complex damage*</u> |  |  |  |
| *Integral*<sup>a</sup> | **9.56** | **0.002** | **2** |
| *t<sub>onset</sub>*<sup>a</sup> | 1.05 | 0.374 | 2 |
| *Duration*<sup>a†</sup> | 1.87 | 0.188 | 2 |
| *Shape*<sup>a†</sup> | 1.74 | 0.209 | 2 |
| <u>*Real herbivory*</u> |  |  |  |
| *t<sub>onset</sub>*<sup>b†</sup> | **4.85** | **0.009** | **4** |
| *Duration*<sup>a*</sup> | **26.72** | **< 0.001** | **4** |
| *Shape*<sup>a†</sup> | **6.91** | **< 0.001** | **4** |

**Supplemental Table 5.** Summary of statistical analyses presented in Supplemental figures 1 and 2. Bold values: *p* < 0.05. a = analysed using one-way ANOVA, b = analysed using Welch’s ANOVA.

| **Response** | ***F*** | ***p-value*** | ***df*** |
| --- | --- | --- | --- |
| <u>*Green leaf volatiles*</u> |  |  |  |
| *t<sub>onset</sub>*<sup>a</sup> | 0.826 | 0.482 | 2 |
| *Duration*<sup>a</sup> | **43.42** | **< 0.001** | **2** |
| *Shape*<sup>a</sup> | **5.43** | **0.045** | **2** |
| <u>*Gene expression*</u> |  |  |  |
| *t<sub>onset</sub>*<sup>b</sup> | **198.42** | **< 0.001** | **3** |
| *Duration*<sup>a</sup> | **54.52** | **< 0.001** | **3** |
| *Shape*<sup>a</sup> | **8.55** | **0.002** | **3** |

## Appendix 1

### Practical Guide to Fitting the Response Model in Python, R, and Excel

This provides a straightforward workflow to apply the response model to experimental time-series data. Each section (Python, R, Excel) can be read independently.

The goal is simple: given a time vector and a response vector, fit the model and extract the derived quantities (shape, duration, total integral).

To apply the model, you only need two data vectors: **time** and **response**. Provide rough starting values or bounds for the four parameters, then run the fitting procedure in your chosen environment (Python, R, or Excel). The steps are the same everywhere:

1. Load or enter your time and response data
2. Set initial parameter guesses or bounds
3. Run the optimiser (Python: DE→SLSQP; R: DEoptim→nloptr; Excel: Solver)
4. Check the fitted curve against your raw data
5. Use the fitted parameters to compute the derived quantities (shape, duration, and total integral)

### Python

#### Define the Model and Fitting Function

This section defines the components needed to run the model:

1. The model, which evaluates the response for any set of parameters.
2. Helper functions that compute the derived quantities (shape, duration, total integral).
3. The fitting function, which estimates the four parameters from data.

The model is a direct implementation of the form described in the manuscript. A gamma-shaped rise and decay captures the response dynamics, while a logistic onset term smooths the beginning of the curve to improve numerical stability. The helper functions compute the analytical quantities defined in the paper.

The main fitting routine fit_response_curve takes three inputs: time , response , and a set of parameter bounds.

These bounds can be estimated visually:

- t_onset : where the rise begins
- t_peak : where the maximum occurs
- t_mean : a point on the decay tail
- R_peak : approximate peak height

The fitting proceeds in two steps. A Differential Evolution search explores the full parameter space without relying on good initial guesses. A constrained SLSQP refinement then improves accuracy while ensuring valid parameter ordering (t_onset < t_peak < t_mean). The result is a dictionary containing the four optimised parameters.

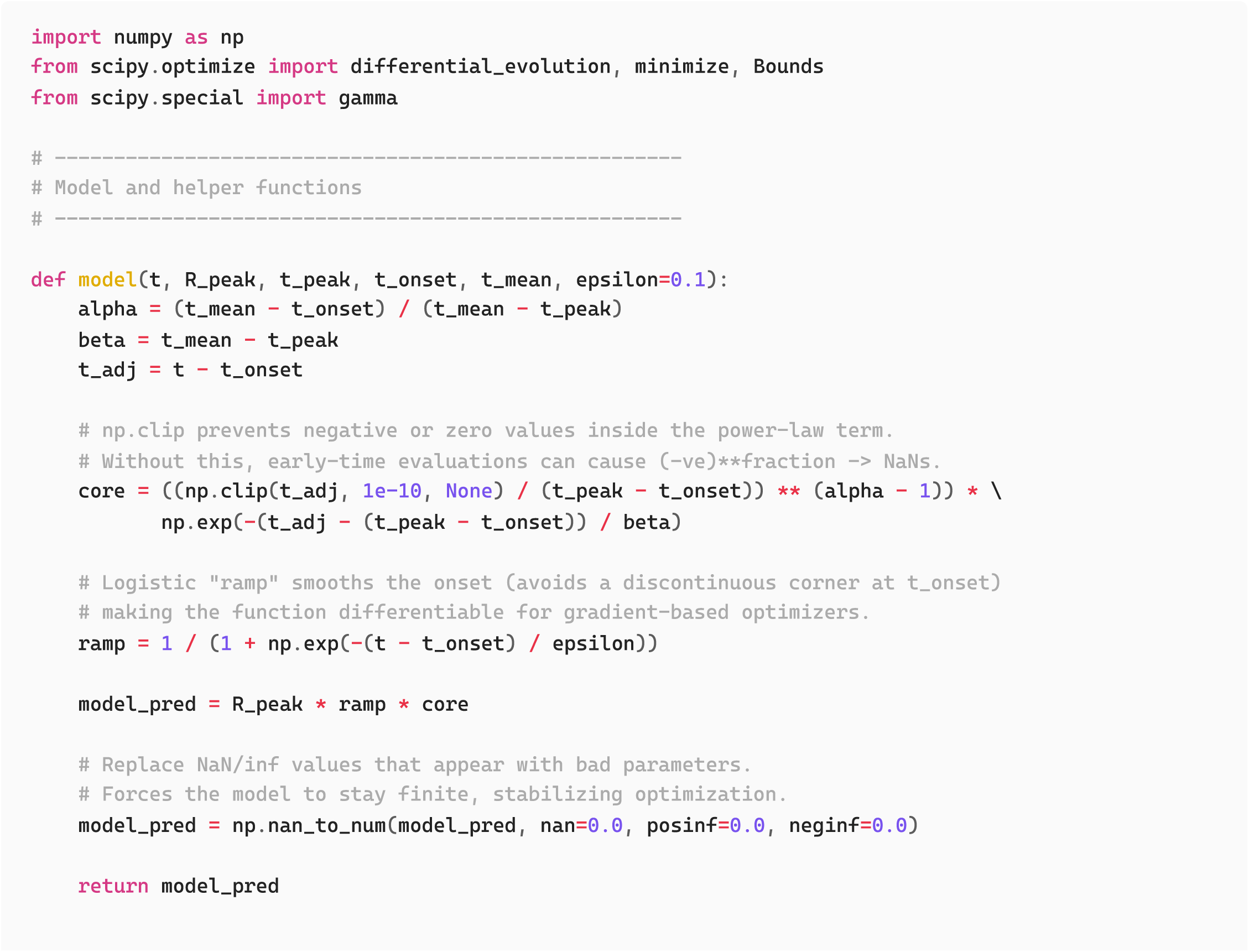
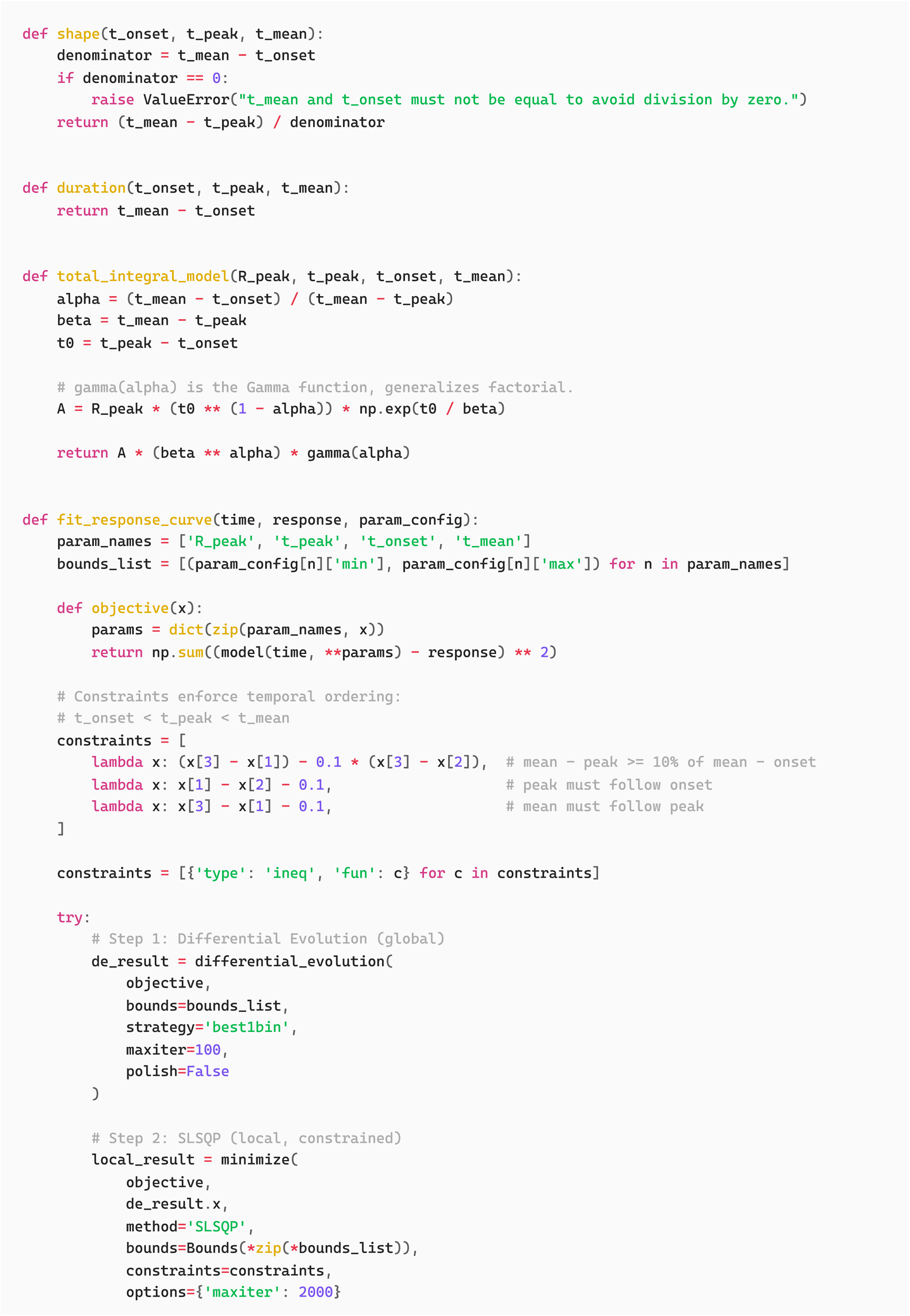
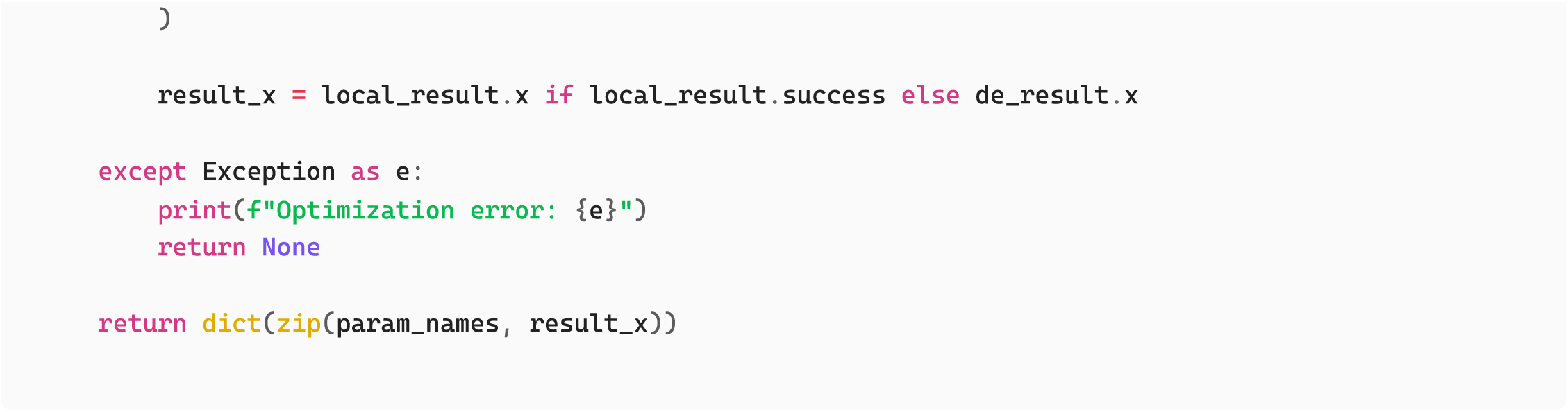

### Fit the model

To run the workflow, provide arrays for time and response, set reasonable bounds, and call fit_response_curve . The fitted parameters can then be used to generate a model prediction and to compute shape, duration, and total integral.

A plot of raw versus fitted data helps assess quality. A good fit should capture the onset, peak position, and overall decay shape. Large systematic deviations usually indicate that bounds were too restrictive or the data deviate from model assumptions.

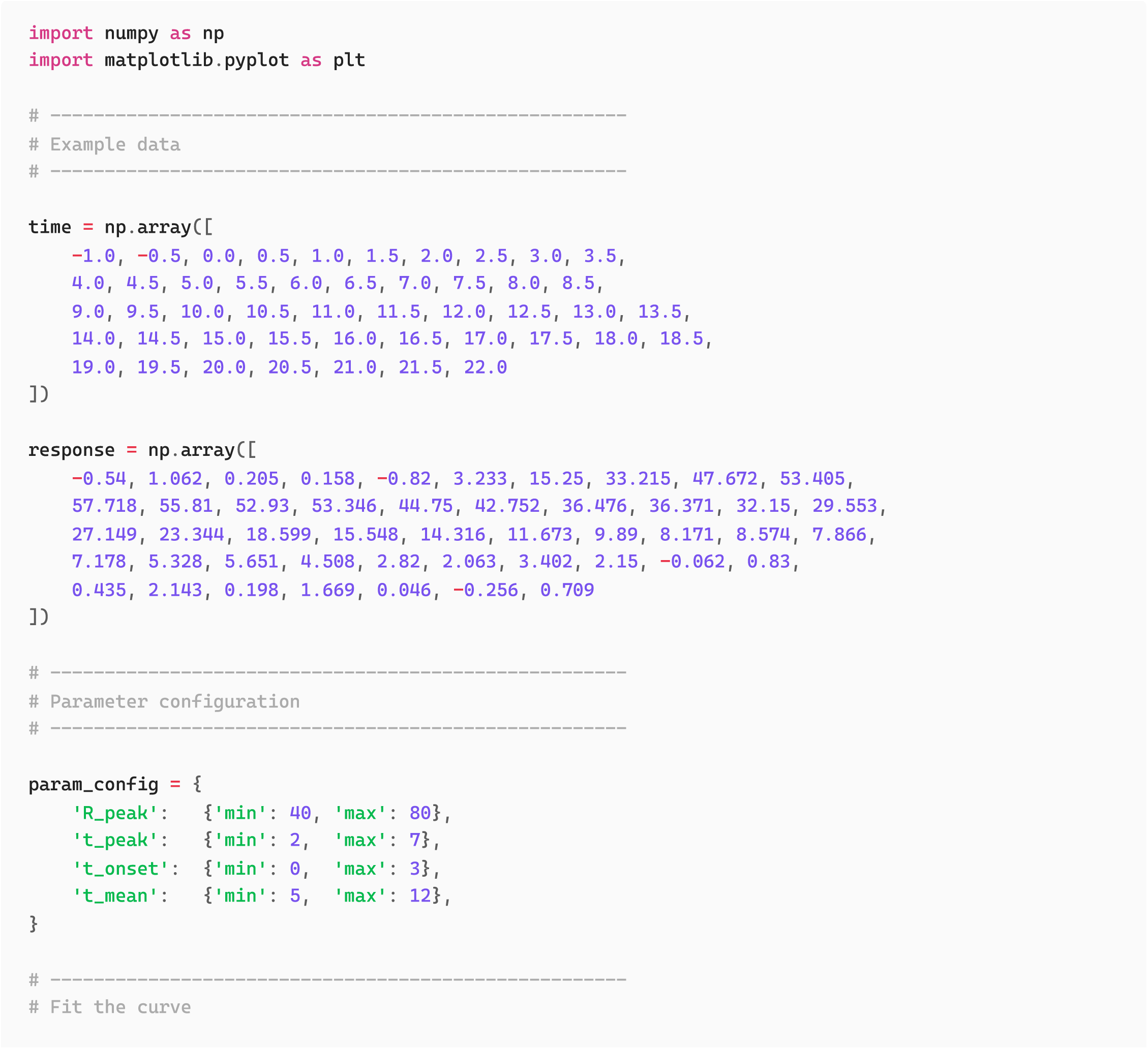
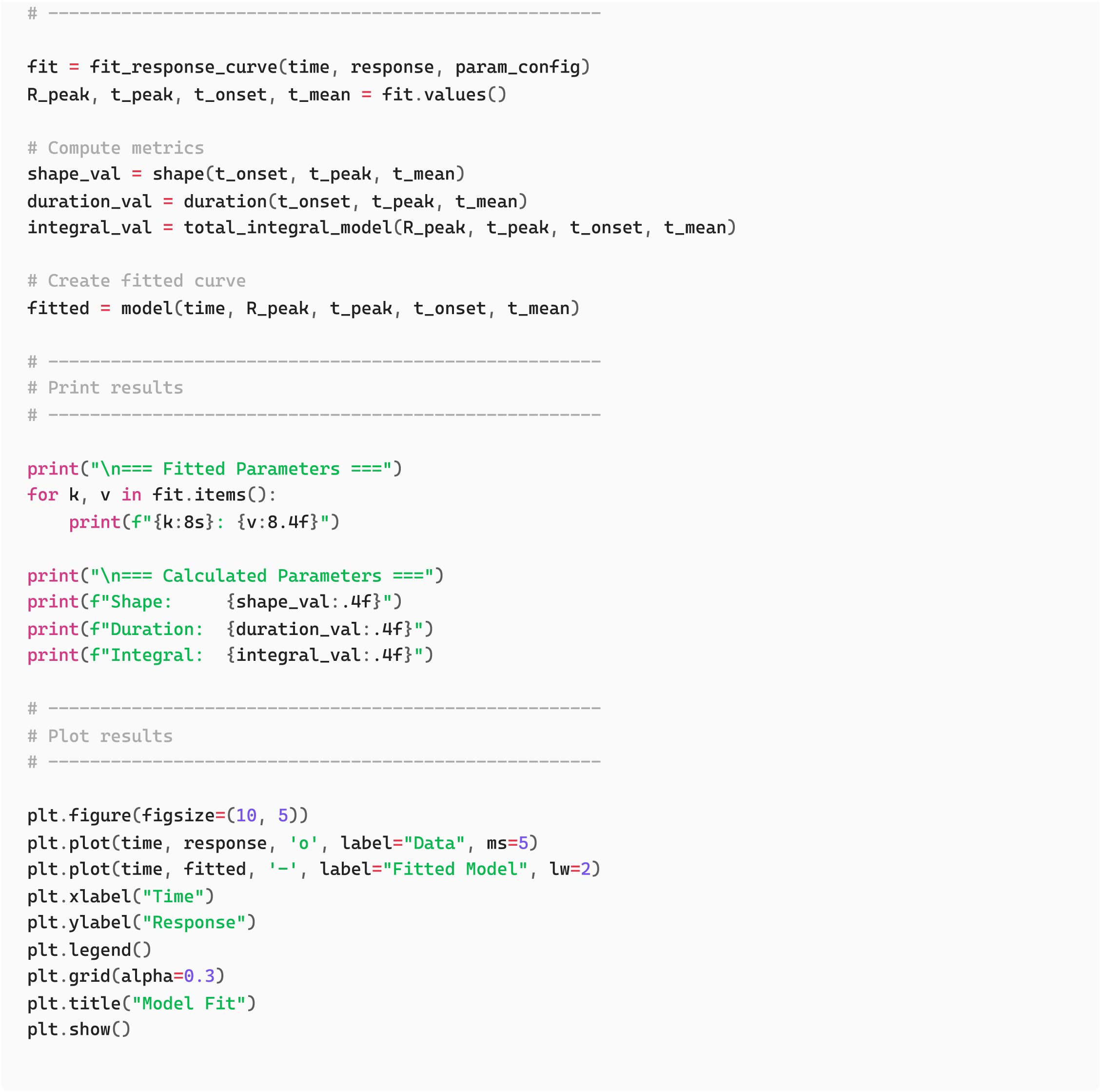

### Output: Python

The printed results show the four fitted parameters and the derived quantities:

- R_peak : fitted peak height
- t_onset , t_peak , t_mean : timing of the main phases
- Shape , Duration , Integral : derived measures of the response

The accompanying plot confirms whether the fitted curve follows the raw data.

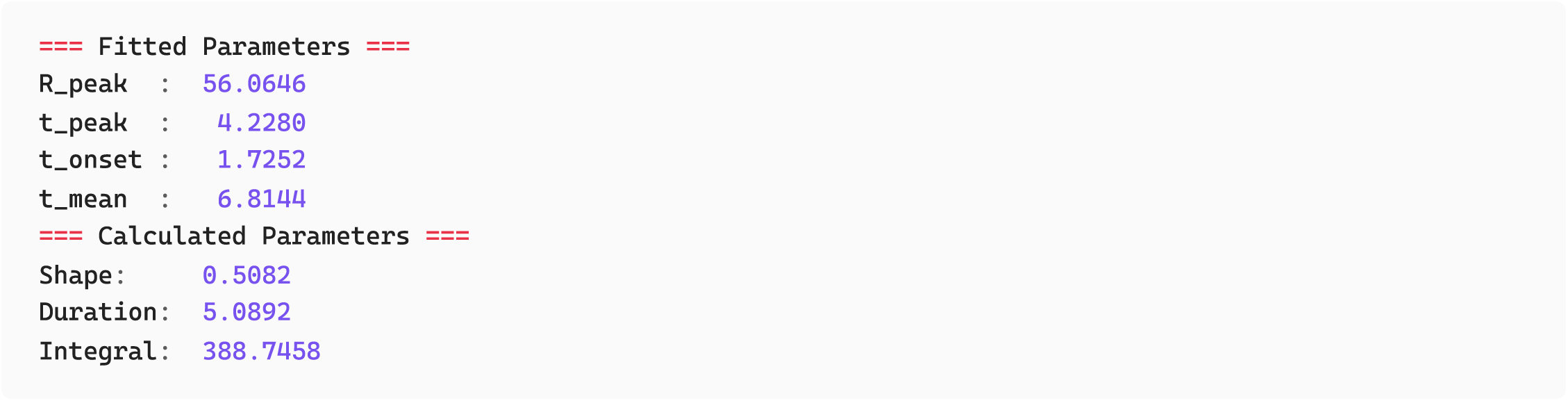

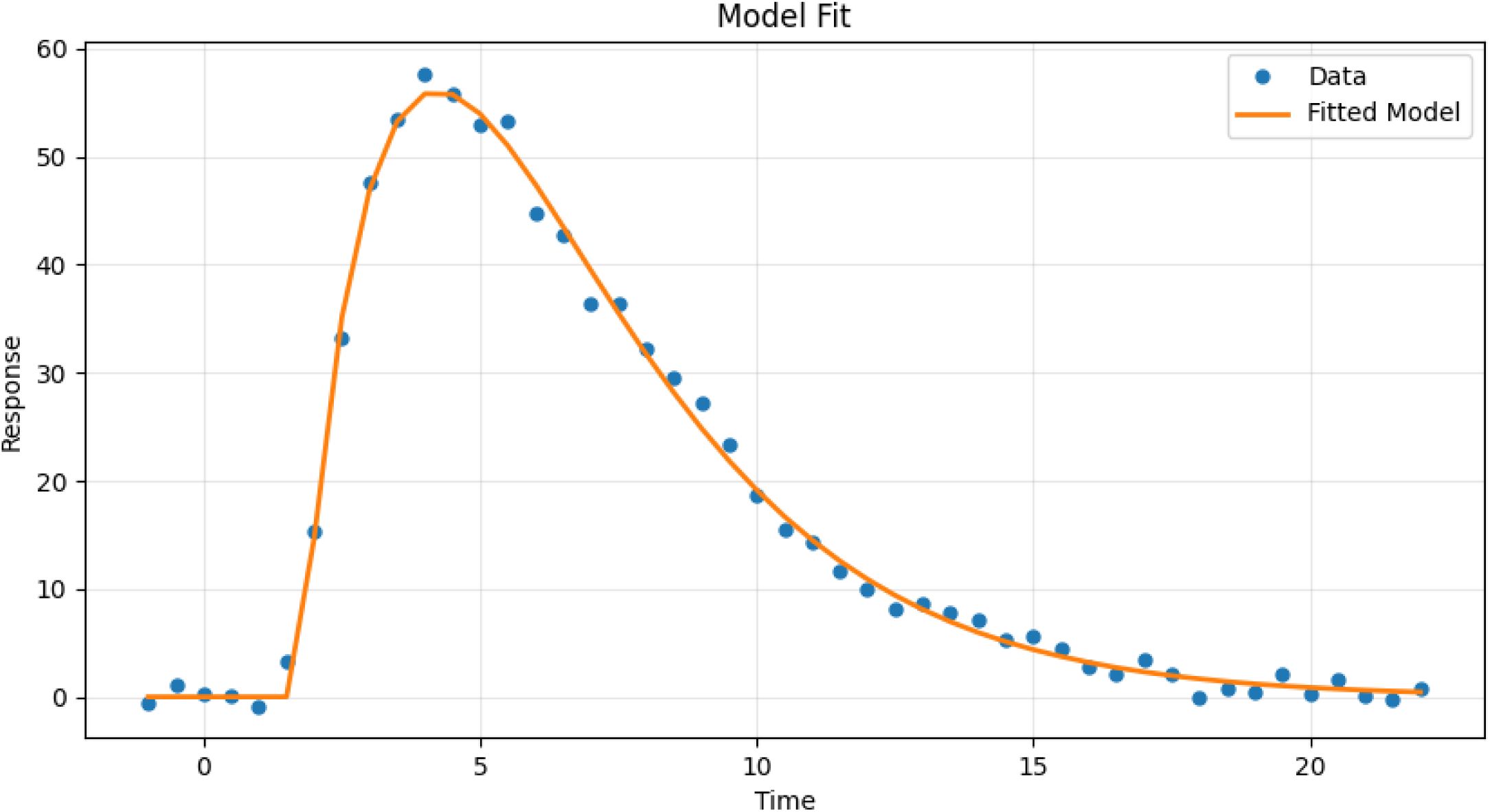

### R: Model and Fitting Functions

This section provides a complete workflow for fitting the response model in R. It includes three components:

1. The model function, which computes the predicted response at each time point.
2. Helper functions that calculate the derived quantities (shape, duration, total integral).
3. A fitting function that estimates the four parameters from experimental data.

The model is a direct implementation of the analytical form described in the manuscript. It combines a gamma-shaped rise and decay with a logistic onset term, which smooths the beginning of the curve and stabilises numerical optimisation. The helper functions use the fitted parameters to compute the analytical descriptors of interest.

The fitting routine fit_response_curve takes three inputs:

- time : numeric vector of time values
- response : numeric vector of observed responses
- param_config : named list giving lower and upper bounds for each parameter

These bounds can be chosen by eye:

- t_onset : near the initial rise
- t_peak : near the maximum
- t_mean : in the decay region
- R_peak : near the observed peak height

The optimisation proceeds in two stages. A DEoptim global search explores the parameter space without relying on good initial guesses. A local refinement using optim (L-BFGS-B) then improves accuracy while keeping parameters within their bounds. Simple inequality checks enforce the required temporal order (t_onset < t_peak < t_mean). The function returns all four fitted parameters in a named list.

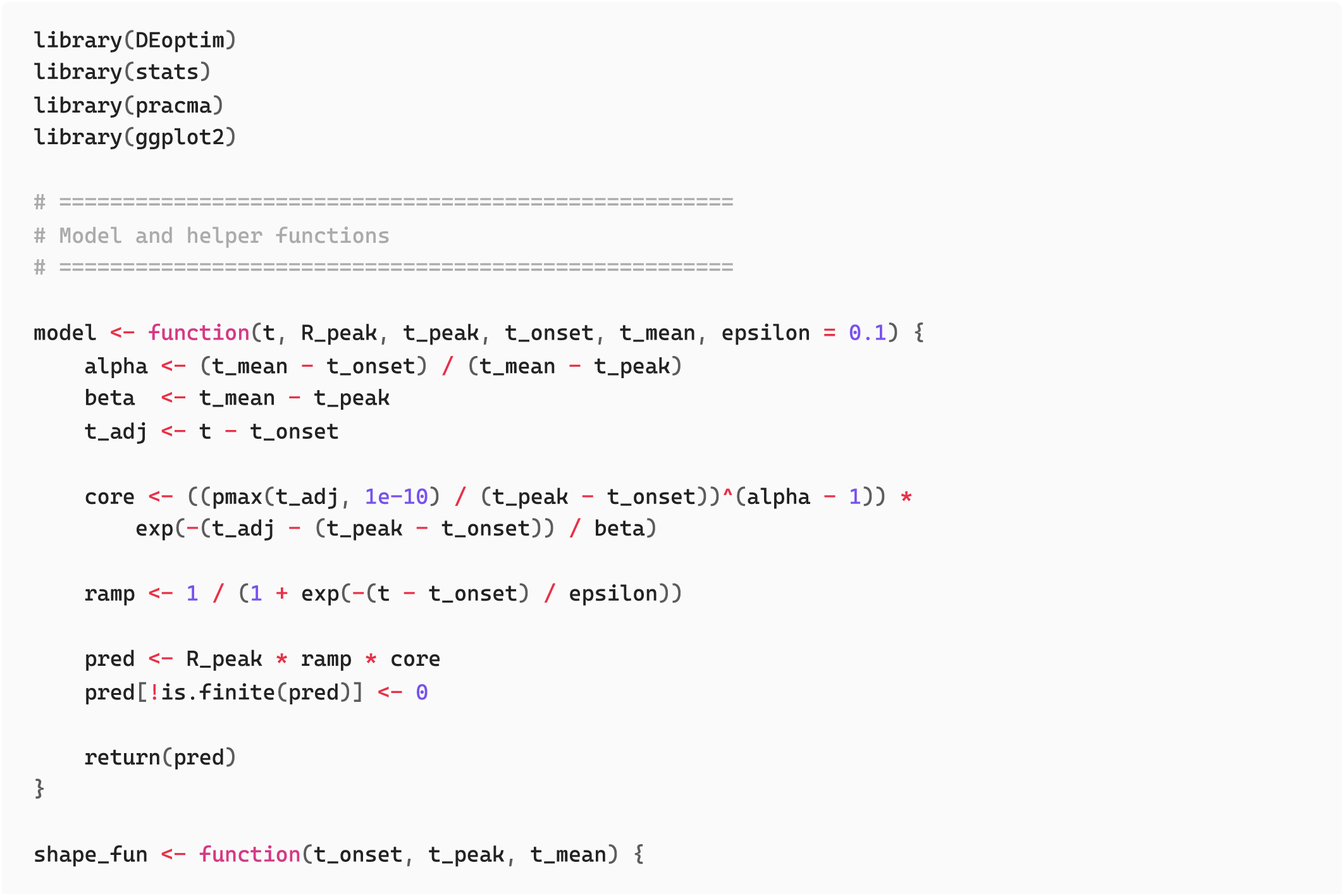
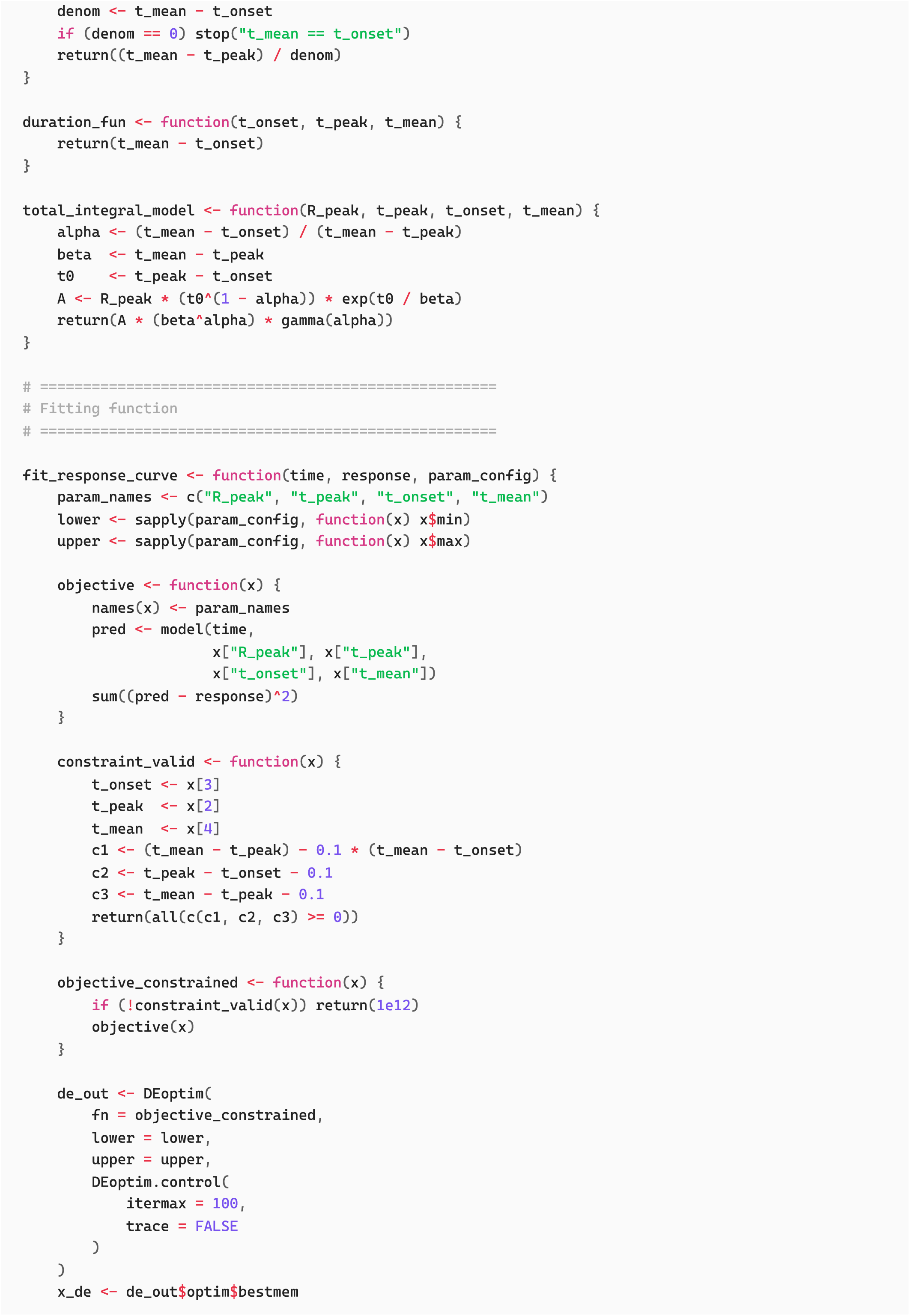
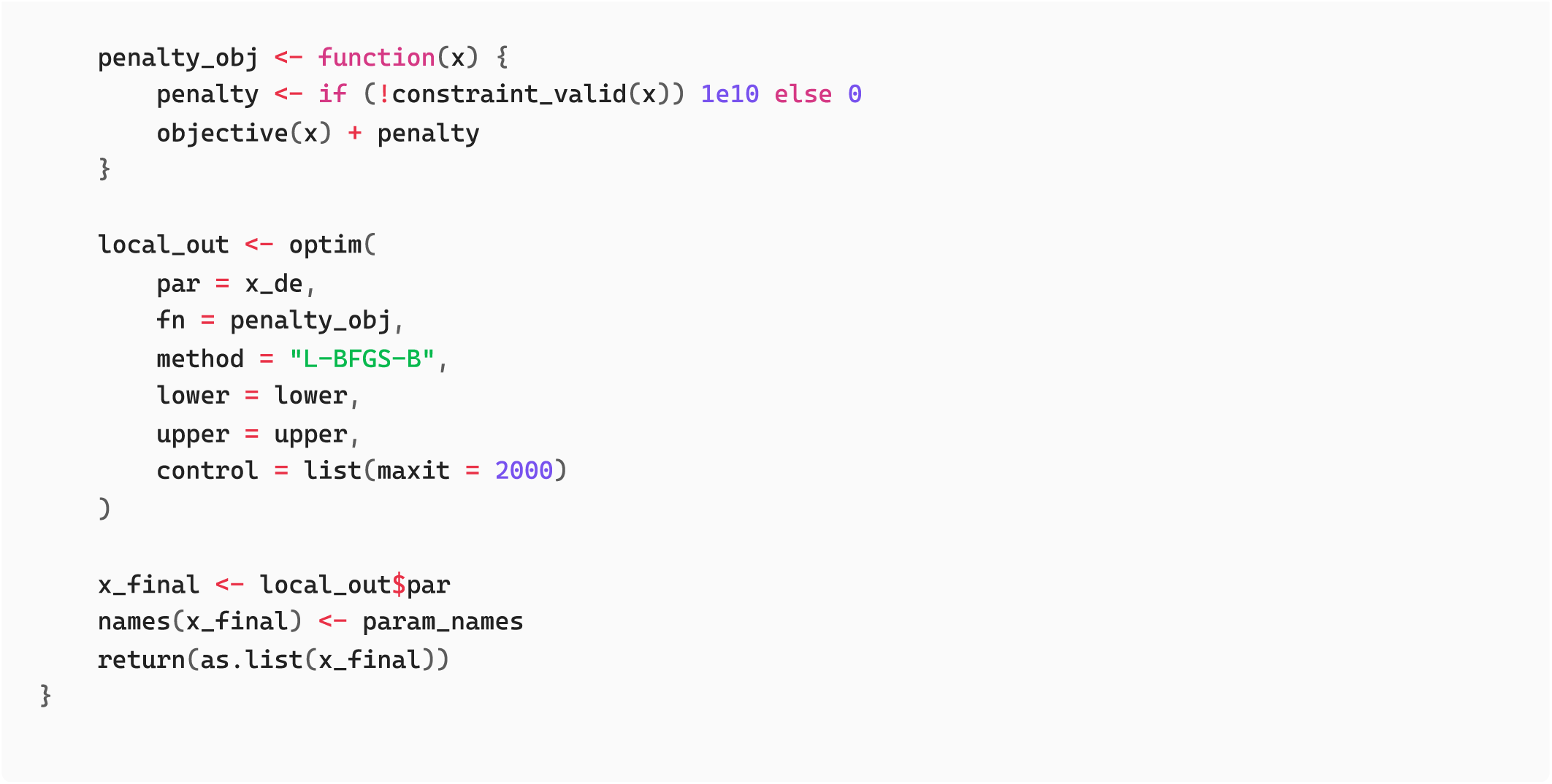

### Usage Example: R

The following example shows the complete workflow: providing time and response vectors, defining parameter bounds, running the fitting routine, and computing the derived quantities. After fitting, the parameters are used both to generate the model prediction and to calculate shape, duration, and integral.

A plot comparing the raw data with the fitted curve provides an immediate check on fit quality. Good fits typically recover the onset, location of the peak, and overall decay structure, though small deviations in noisy regions are expected.

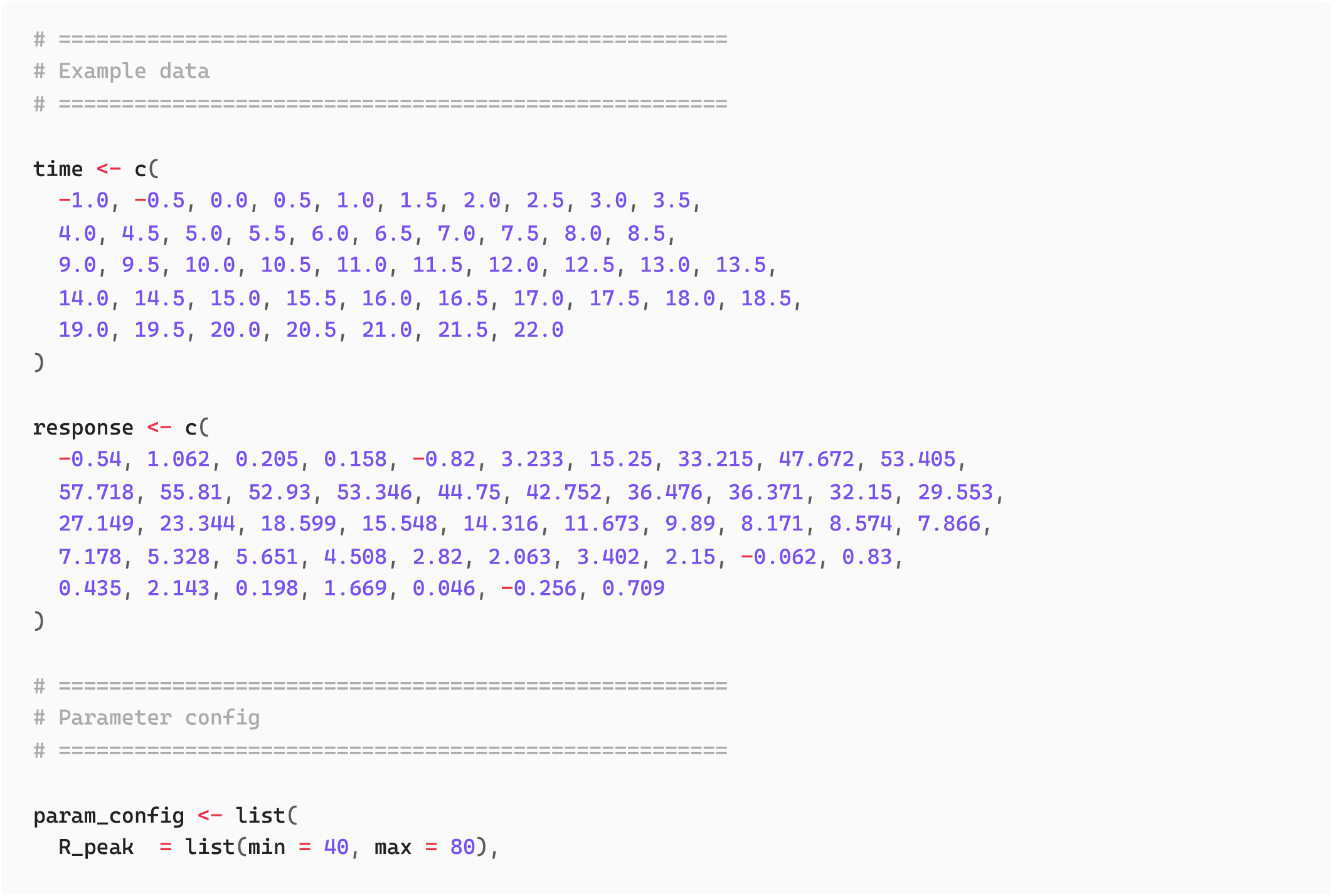
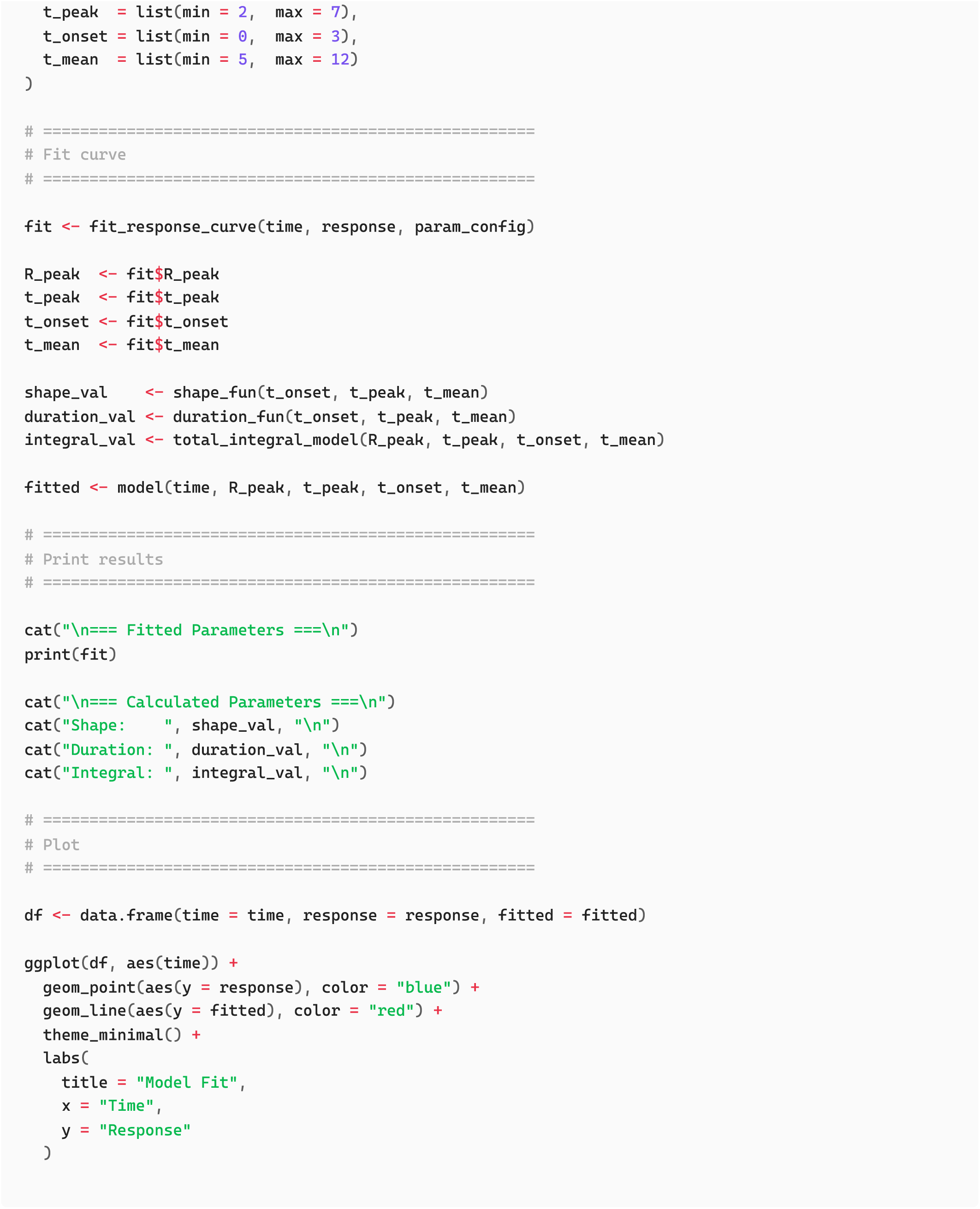

### Output: R

The fitted parameters and derived quantities are printed in a straightforward format:

- R_peak : fitted maximum response
- t_onset , t_peak , t_mean : timing of the response phases
- Shape , Duration , Integral : analytical quantities derived from the fitted model

The plot confirms visually whether the fitted curve captures the key features of the observed data.

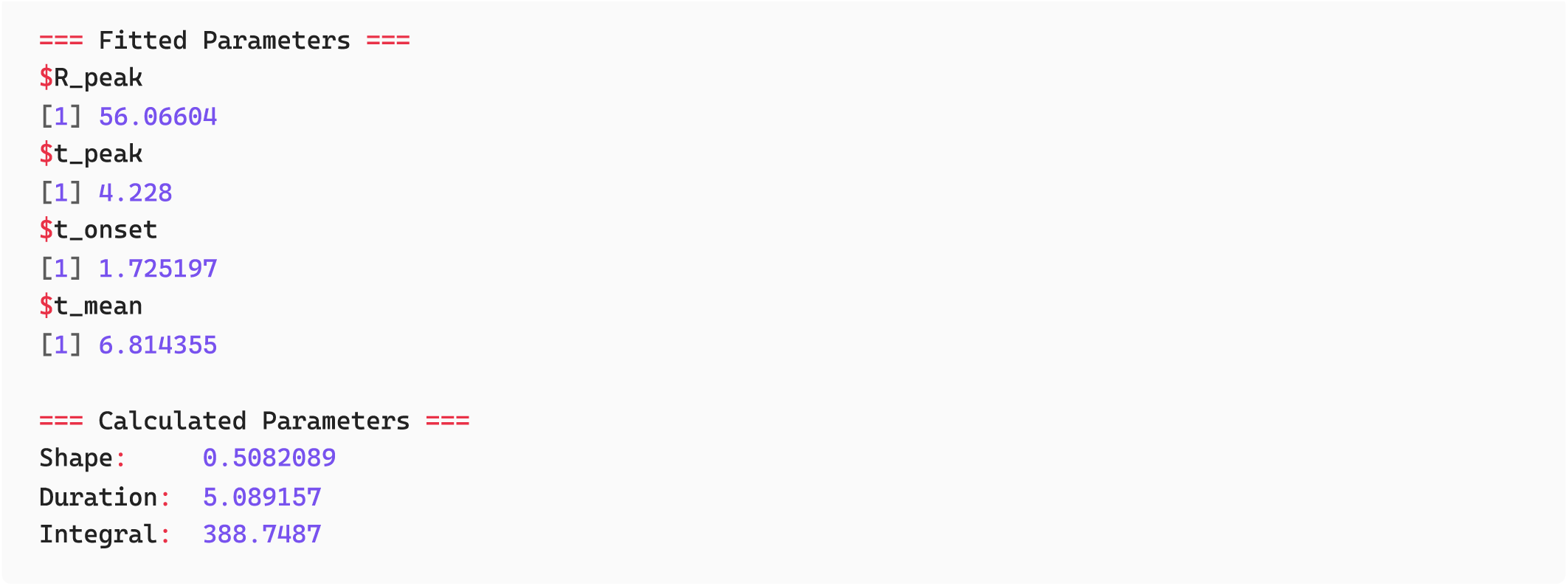

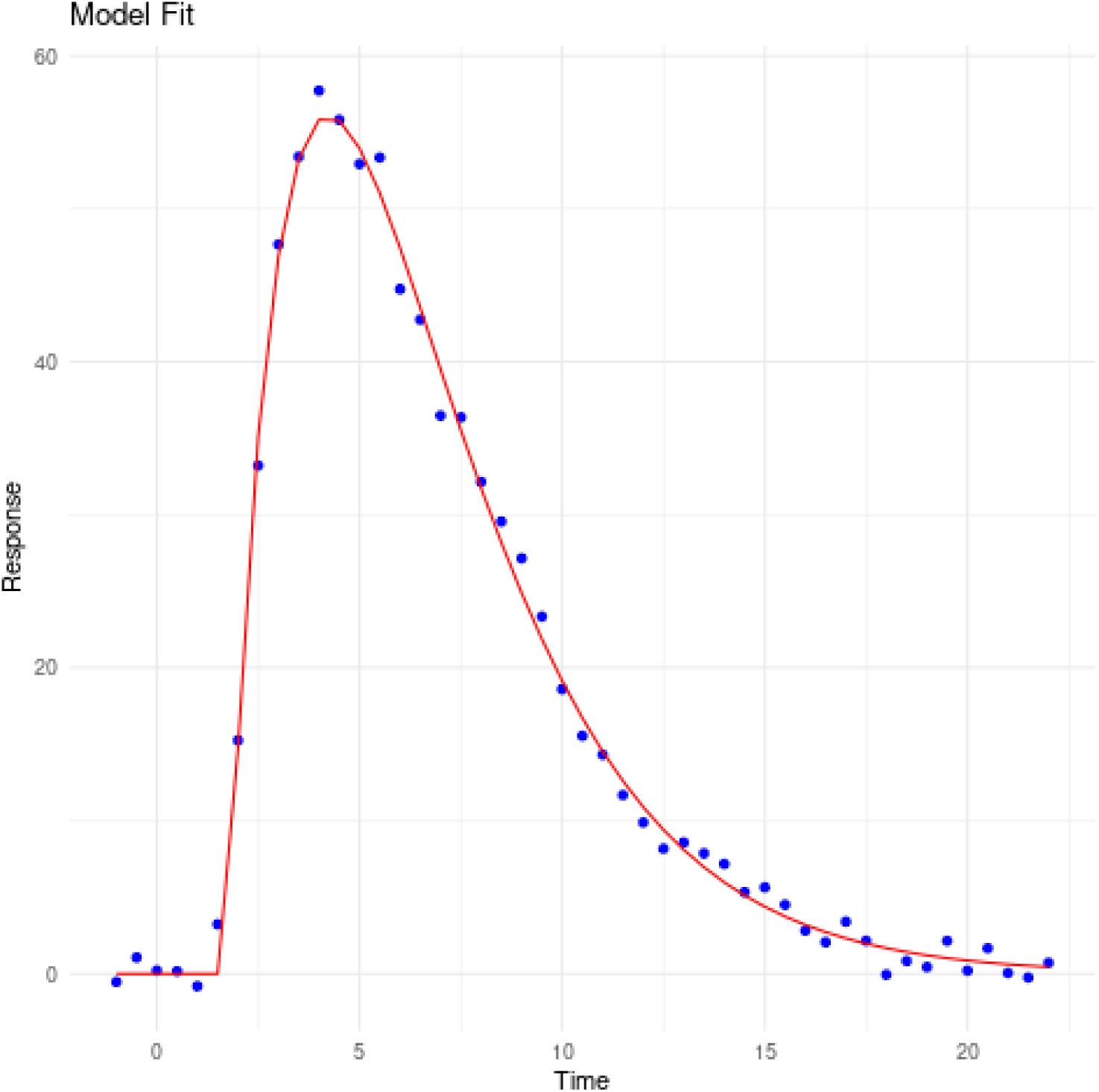

### Excel: Model and Fitting Functions

Excel provides a simple, visual way to use the model. The full workflow is:

1. Enter your time and response data
2. Enter initial parameter guesses and provide min/max bounds for each parameter
3. Compute the model prediction
4. Compute squared errors and SSE
5. Use Solver to optimise the parameters
6. Compute the derived quantities (shape, duration, integral)

#### Step 1

Place your measurements into two columns:

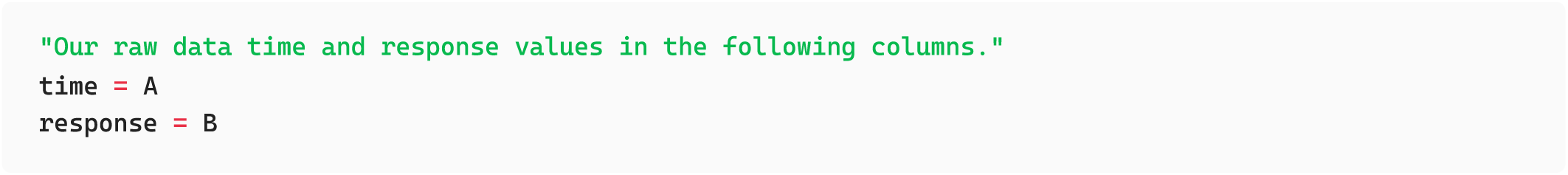

#### Step 2

Reserve cells for the four parameters and their bounds, entering rough starting velues:

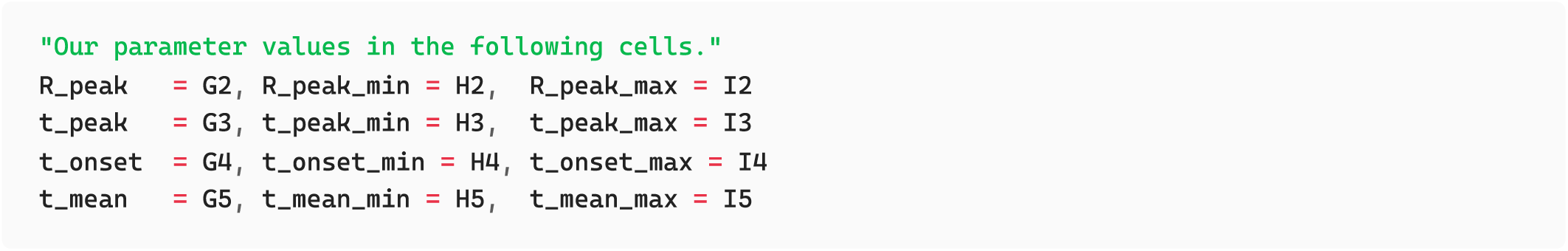

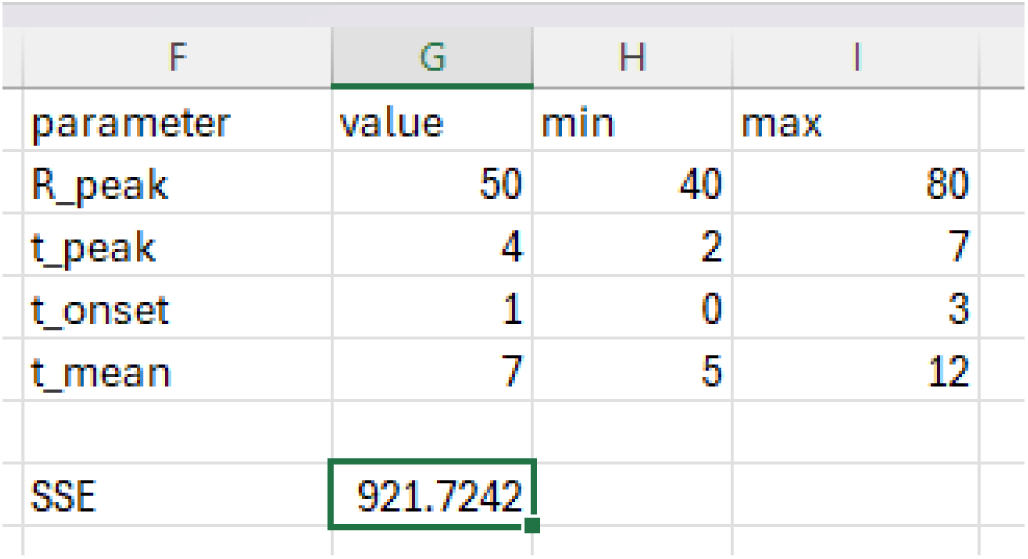

#### Step 3

In **C2**, enter the model formula and drag down:

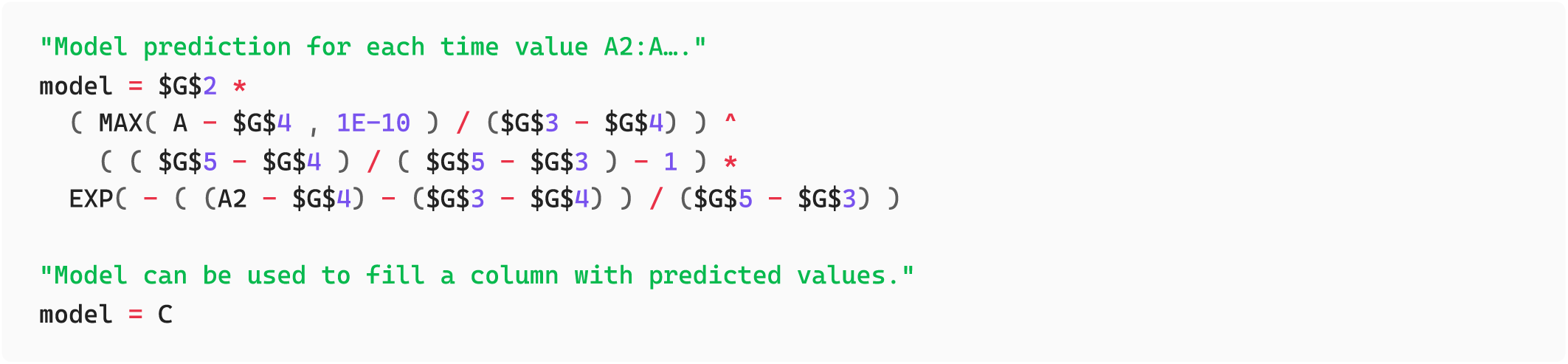

Column **C** now contains the predicted values.

#### Step 4

In order to fit the model, we first need a way to measure how far the current parameter guesses are from the actual data. We do this by calculating the **squared error** between the observed response and the model prediction at each time point.

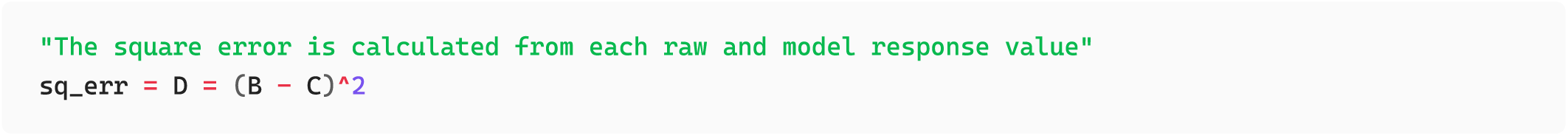

Drag this down to fill the column.

To combine all point-wise errors into a single value that Solver can minimise, compute the sum of squared errors (SSE) in G10:

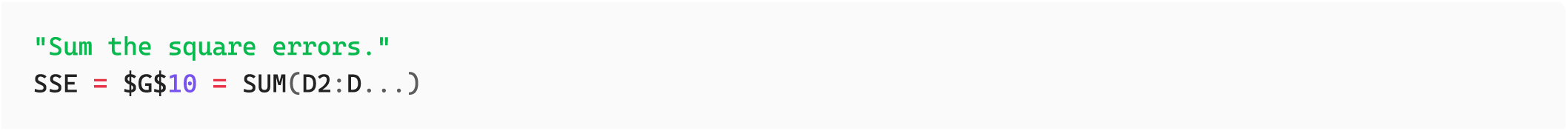

The SSE reflects how well the current parameters match the data: lower values indicate a better fit. Solver will adjust the parameters to minimise this value.

#### Step 5

Open **Data → Solver** and configure the following:

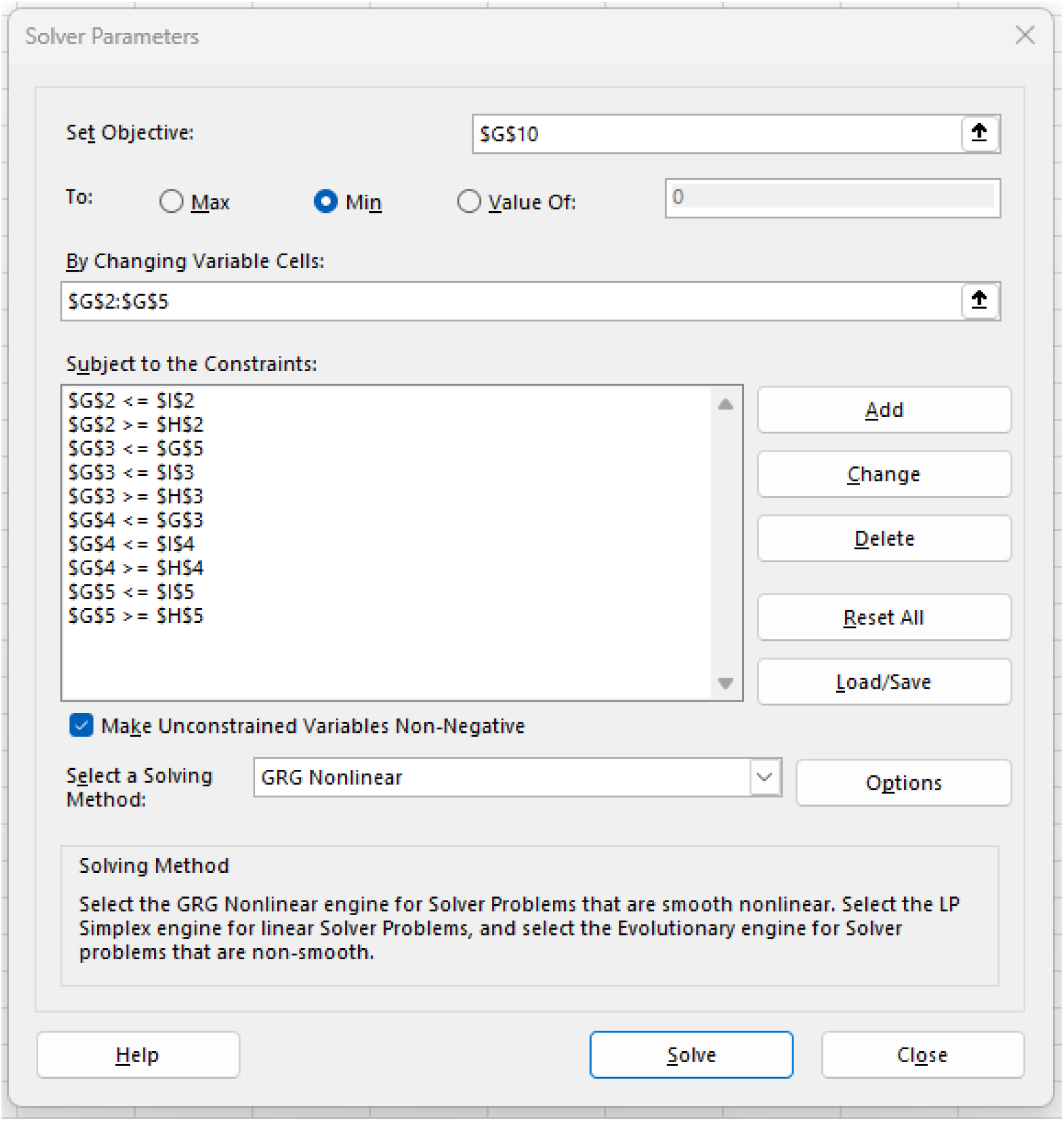

### Objective

- Set Objective :
- To : Min Variable cells

#### Constraints

Add both parameter bounds and temporal ordering:

Subject to the Constraints:

G2 >= H2 and G2 <= I2

G3 >= H3 and G3 <= I3

G4 >= H4 and G4 <= I4

G5 >= H5 and G5 <= I5

#### Temporal ordering

- G4 < G3 (onset before peak)
- G3 < G5 (peak before mean)

### Method

- Select a Solving Method : GRG Nonlinear

Click Solve .

If successful, Solver updates cells with the best-fitting parameters.

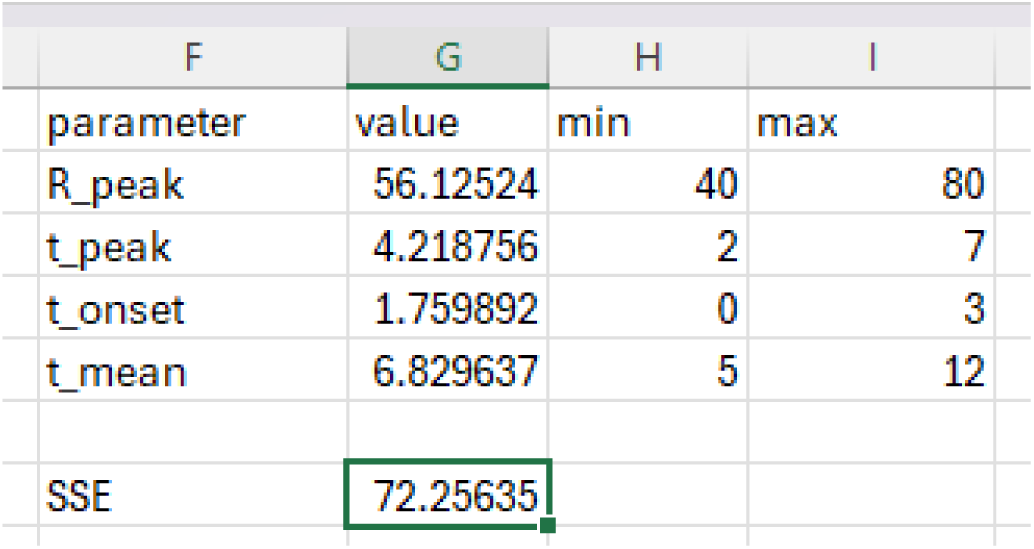

#### Step 6

Once Solver converges, compute the analytical quantities that characterise the fitted response. In any convenient column or cells:

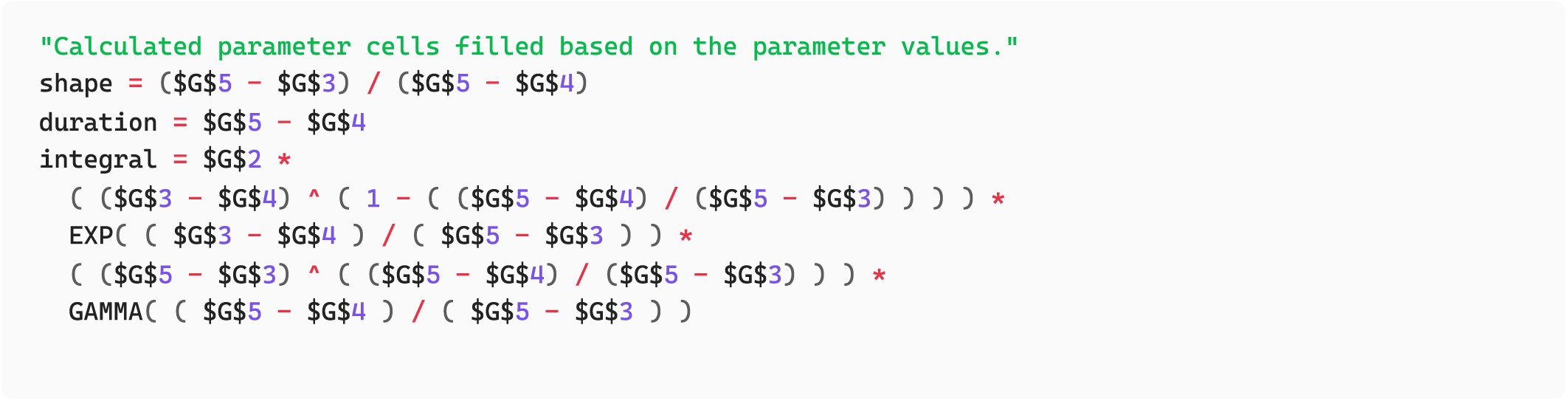

These update automatically if the parameter cells change.

##### Output: Excel

After running Solver, the fitted parameter values appear in the parameter cells, and the SSE decreases accordingly. The model prediction column updates automatically, as do the derived quantities (shape, duration, integral). These values provide a direct summary of the fitted response.

A plot comparing the observed data with the model prediction gives a visual check of fit quality. A successful fit will show the model curve following the rise, peak, and decay of the measured response, as illustrated in the example screenshot.

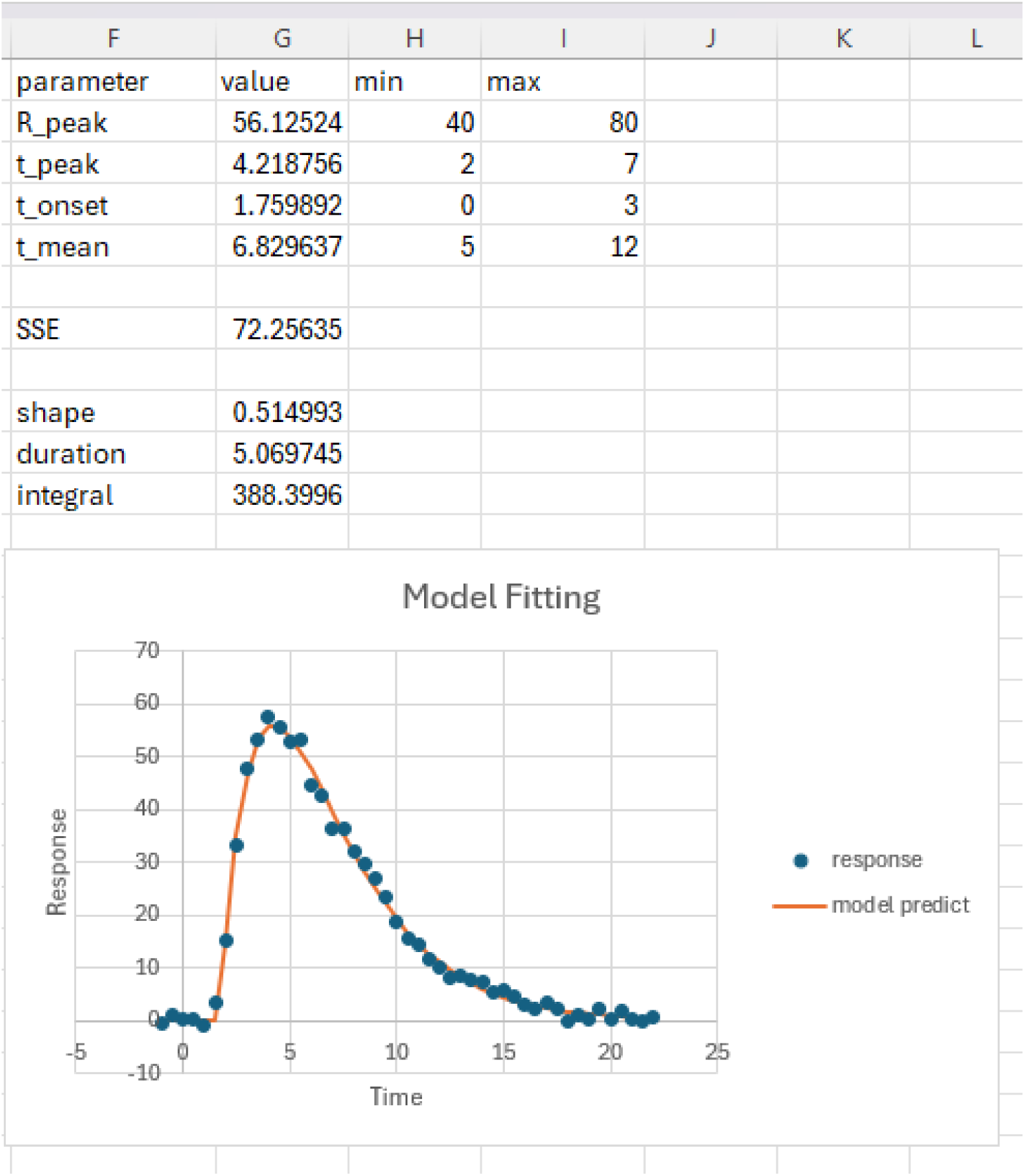

## References

1. M. Erb, N. Veyrat, C. A. M. Robert, H. Xu, M. Frey, J. Ton, T. C. J. Turlings, Indole is an essential herbivore-induced volatile priming signal in maize. Nature Communications 6, 6273 (2015).

2. Y.-Q. Gao, P. Jimenez-Sandoval, S. Tiwari, S. Stolz, J. Wang, G. Glauser, J. Santiago, E. E. Farmer, Ricca’s factors as mobile proteinaceous effectors of electrical signaling. Cell 186, 1337–1351.e20 (2023).

3. H. P. Lu, L. Xun, X. S. Xie, Single-Molecule Enzymatic Dynamics. Science 282, 1877–1882 (1998).

4. M. Toyota, D. Spencer, S. Sawai-Toyota, W. Jiaqi, T. Zhang, A. J. Koo, G. A. Howe, S. Gilroy, Glutamate triggers long-distance, calcium-based plant defense signaling. Science 361, 1112–1115 (2018).

5. L. Yang, V. Pathiranage, S. Zhou, X. Sun, H. Zhang, C. Lai, C. Gu, F. V. Subach, M. Drobizhev, A. R. Walker, K. D. Piatkevich, Sensitive red fluorescent indicators for real-time visualization of potassium ion dynamics in vivo. PLOS Biology 23, e3002993 (2025).

6. M. C.-Y. Ang, J. M. Saju, T. K. Porter, S. Mohaideen, S. Sarangapani, D. T. Khong, S. Wang, J. Cui, S. I. Loh, G. P. Singh, N.-H. Chua, M. S. Strano, R. Sarojam, Decoding early stress signaling waves in living plants using nanosensor multiplexing. Nature Communications 15, 2943 (2024).

7. P. Coatsworth, L. Gonzalez-Macia, A. S. P. Collins, T. Bozkurt, F. Güder, Continuous monitoring of chemical signals in plants under stress. Nature Reviews Chemistry 7, 7–25 (2023).

8. R. Grote, M. Sharma, A. Ghirardo, J.-P. Schnitzler, A New Modeling Approach for Estimating Abiotic and Biotic Stress-Induced de novo Emissions of Biogenic Volatile Organic Compounds From Plants. Frontiers in Forests and Global Change **Volume** 2-2019 (2019).

9. L. Oyarte Galvez, C. Bisot, P. Bourrianne, R. Cargill, M. Klein, M. van Son, J. van Krugten, V. Caldas, T. Clerc, K.-K. Lin, F. Kahane, S. van Staalduine, J. D. Stewart, V. Terry, B. Turcu, S. van Otterdijk, A. Babu, M. Kamp, M. Seynen, B. Steenbeek, J. Zomerdijk, E. Tutucci, M. Sheldrake, C. Godin, V. Kokkoris, H. A. Stone, E. T. Kiers, T. S. Shimizu, A travelling-wave strategy for plant–fungal trade. Nature 639, 172–180 (2025).

10. J. M. Waterman, T. M. Cofer, L. Wang, G. Glauser, M. Erb, High-resolution kinetics of herbivore-induced plant volatile transfer reveal clocked response patterns in neighboring plants. eLife 12, RP89855 (2024).

11. J. Zhang, H. Yin, J. Zhang, G. Yang, J. Qin, L. He, Real-time mental stress detection using multimodality expressions with a deep learning framework. Frontiers in Neuroscience **Volume** 16–2022 (2022).

12. Ü. Niinemets, A. Kännaste, L. Copolovici, Quantitative patterns between plant volatile emissions induced by biotic stresses and the degree of damage. Frontiers in Plant Science **Volume** 4-2013 (2013).

13. J. Reyes-Silveyra, A. R. Mikler, Modeling immune response and its effect on infectious disease outbreak dynamics. Theoretical Biology and Medical Modelling 13, 10 (2016).

14. V. S. Pan, E. Ghosh, P. J. Ode, W. C. Wetzel, K. J. Gilbert, I. S. Pearse, Large Differences in Herbivore Performance Emerge From Simple Herbivore Behaviours and Fine-Scale Spatial Heterogeneity in Phytochemistry. Ecology Letters 28, e70044 (2025).

15. I. S. Pearse, R. Paul, P. J. Ode, Variation in Plant Defense Suppresses Herbivore Performance. Current Biology 28, 1981–1986.e2 (2018).

16. W. C. Wetzel, H. M. Kharouba, M. Robinson, M. Holyoak, R. Karban, Variability in plant nutrients reduces insect herbivore performance. Nature 539, 425–427 (2016).

17. T. Branco, B. A. Clark, M. Häusser, Dendritic Discrimination of Temporal Input Sequences in Cortical Neurons. Science 329, 1671–1675 (2010).

18. B. Grothe, M. Pecka, D. McAlpine, Mechanisms of Sound Localization in Mammals. Physiological Reviews 90, 983–1012 (2010).

19. J. M. Köhler, Vaccination, Immunity and Breakthrough: Quantitative Effects in Individual Immune Responses Illustrated by a Simple Kinetic Model. Applied Sciences 12 (2022).

20. P. A. Gagliardi, O. Pertz, The mitogen-activated protein kinase network, wired to dynamically function at multiple scales. Current Opinion in Cell Biology 88, 102368 (2024).

21. I. Kinjyo, J. Qin, S.-Y. Tan, C. J. Wellard, P. Mrass, W. Ritchie, A. Doi, L. L. Cavanagh, M. Tomura, A. Sakaue-Sawano, O. Kanagawa, A. Miyawaki, P. D. Hodgkin, W. Weninger, Real-time tracking of cell cycle progression during CD8+ effector and memory T-cell differentiation. Nature Communications 6, 6301 (2015).

22. A. Stirbet, D. Lazár, Y. Guo, G. Govindjee, Photosynthesis: basics, history and modelling. Annals of Botany 126, 511–537 (2020).

23. J. Wu, I. T. Baldwin, Herbivory-induced signalling in plants: perception and action. Plant, Cell & Environment 32, 1161–1174 (2009).

24. X. Yu, B. Feng, P. He, L. Shan, From Chaos to Harmony: Responses and Signaling upon Microbial Pattern Recognition, Annual Review of Phytopathology. 55 (2017)pp. 109–137.

25. J. M. S. Zegers, I. Irisarri, S. de Vries, J. de Vries, Evolving circuitries in plant signaling cascades. Journal of Cell Science 137, jcs261712 (2024).

26. A. K. Block, C. T. Hunter, C. Rering, S. A. Christensen, R. L. Meagher, Contrasting insect attraction and herbivore-induced plant volatile production in maize. Planta 248, 105–116 (2018).

27. R. Moreno-Sánchez, E. Saavedra, S. Rodríguez-Enríquez, V. Olín-Sandoval, Metabolic Control Analysis: A Tool for Designing Strategies to Manipulate Metabolic Pathways. BioMed Research International 2008, 597913 (2008).

28. S. A. R. Mousavi, A. Chauvin, F. Pascaud, S. Kellenberger, E. E. Farmer, GLUTAMATE RECEPTOR-LIKE genes mediate leaf-to-leaf wound signalling. Nature 500, 422–426 (2013).

29. Y. Blum, J. Mikelson, M. Dobrzyński, H. Ryu, M. Jacques, N. L. Jeon, M. Khammash, O. Pertz, Temporal perturbation of ERK dynamics reveals network architecture of FGF2/MAPK signaling. Molecular Systems Biology 15, e8947 (2019).

30. C. Dessauges, J. Mikelson, M. Dobrzyński, M. Jacques, A. Frismantiene, P. A. Gagliardi, M. Khammash, O. Pertz, Optogenetic actuator – ERK biosensor circuits identify MAPK network nodes that shape ERK dynamics. Molecular Systems Biology 18, e10670 (2022).

31. T. K. Porter, M. N. Heinz, D. J. Lundberg, A. M. Brooks, T. T. S. Lew, K. S. Silmore, V. B. Koman, M. C.-Y. Ang, D. T. Khong, G. P. Singh, J. W. Swan, R. Sarojam, N.-H. Chua, M. S. Strano, A theory of mechanical stress-induced H2O2 signaling waveforms in Planta. Journal of Mathematical Biology 86, 11 (2022).

32. T. K. Porter, G. Sánchez-Velázquez, M. S. Strano, The Role of Basal H2O2 Concentration in ROS Stress Signaling Waveforms In Planta. *ACS Agric*. Sci. Technol. 5, 1434–1441 (2025).

33. P. A. Saa, L. K. Nielsen, Construction of feasible and accurate kinetic models of metabolism: A Bayesian approach. Scientific Reports 6, 29635 (2016).

34. L. M. Tran, M. L. Rizk, J. C. Liao, Ensemble Modeling of Metabolic Networks. Biophysical Journal 95, 5606–5617 (2008).

35. A. Khodayari, C. D. Maranas, A genome-scale Escherichia coli kinetic metabolic model k-ecoli457 satisfying flux data for multiple mutant strains. Nature Communications 7, 13806 (2016).

36. H. Link, D. Christodoulou, U. Sauer, Advancing metabolic models with kinetic information. Current Opinion in Biotechnology 29, 8–14 (2014).

37. R. N. Gutenkunst, J. J. Waterfall, F. P. Casey, K. S. Brown, C. R. Myers, J. P. Sethna, Universally Sloppy Parameter Sensitivities in Systems Biology Models. PLOS Computational Biology 3, e189 (2007).

38. M. E. Bergman, R. W. J. Kortbeek, M. Gutensohn, N. Dudareva, Plant terpenoid biosynthetic network and its multiple layers of regulation. Progress in Lipid Research 95, 101287 (2024).

39. J.-K. Weng, The evolutionary paths towards complexity: a metabolic perspective. New Phytologist 201, 1141–1149 (2014).

40. J. P. Torres, Z. Lin, J. M. Winter, P. J. Krug, E. W. Schmidt, Animal biosynthesis of complex polyketides in a photosynthetic partnership. Nature Communications 11, 2882 (2020).

41. J. R. Stinchcombe, M. Kirkpatrick, Genetics and evolution of function-valued traits: understanding environmentally responsive phenotypes. Trends in Ecology & Evolution 27, 637–647 (2012).

42. Izem Rima, Kingsolver Joel G., Brodie Edmund D., Losos Jonathan B., Variation in Continuous Reaction Norms: Quantifying Directions of Biological Interest. The American Naturalist 166, 277–289 (2005).

43. M. E. De Obaldia, T. Morita, L. C. Dedmon, D. J. Boehmler, C. S. Jiang, E. V. Zeledon, J. R. Cross, L. B. Vosshall, Differential mosquito attraction to humans is associated with skin-derived carboxylic acid levels. Cell 185, 4099–4116.e13 (2022).

44. R. Escobar-Bravo, P.-A. Lin, J. M. Waterman, M. Erb, Dynamic environmental interactions shaped by vegetative plant volatiles. Nat. Prod. Rep. 40, 840–865 (2023).

45. A. M. Angioy, A. Desogus, I. T. Barbarossa, P. Anderson, B. S. Hansson, Extreme Sensitivity in an Olfactory System. Chemical Senses 28, 279–284 (2003).

46. H. Guo, D. P. Smith, Time-Dependent Odorant Sensitivity Modulation in Insects. Insects 13 (2022).

47. P. Zu, K. Boege, E. del-Val, M. C. Schuman, P. C. Stevenson, A. Zaldivar-Riverón, S. Saavedra, Information arms race explains plant-herbivore chemical communication in ecological communities. Science 368, 1377–1381 (2020).

48. Y. Qin, L. Wang, D. Zhong, Dynamics and mechanism of ultrafast water–protein interactions. Proceedings of the National Academy of Sciences 113, 8424–8429 (2016).

49. M. Shibata, H. Nishimasu, N. Kodera, S. Hirano, T. Ando, T. Uchihashi, O. Nureki, Real-space and real-time dynamics of CRISPR-Cas9 visualized by high-speed atomic force microscopy. Nature Communications 8, 1430 (2017).

50. E. T. Wurtzel, T. M. Kutchan, Plant metabolism, the diverse chemistry set of the future. Science 353, 1232–1236 (2016).

51. Y. T. Cheng, C. A. Thireault, L. Zhang, B. C. Paasch, R. Sohrabi, S. Y. He, Roles of microbiota in autoimmunity in Arabidopsis leaves. Nature Plants 10, 1363–1376 (2024).

52. J.-H. Park, R. Halitschke, H. B. Kim, I. T. Baldwin, K. A. Feldmann, R. Feyereisen, A knock-out mutation in allene oxide synthase results in male sterility and defective wound signal transduction in Arabidopsis due to a block in jasmonic acid biosynthesis. The Plant Journal 31, 1–12 (2002).

53. T. C. J. Turlings, U. B. Lengwiler, M. L. Bernasconi, D. Wechsler, Timing of induced volatile emissions in maize seedlings. Planta 207, 146–152 (1998).

54. B. Loubet, P. Buysse, L. Gonzaga-Gomez, F. Lafouge, R. Ciuraru, C. Decuq, J. Kammer, S. Bsaibes, C. Boissard, B. Durand, J.-C. Gueudet, O. Fanucci, O. Zurfluh, L. Abis, N. Zannoni, F. Truong, D. Baisnée, R. Sarda-Estève, M. Staudt, V. Gros, Volatile organic compound fluxes over a winter wheat field by PTR-Qi-TOF-MS and eddy covariance. Atmospheric Chemistry and Physics 22, 2817–2842 (2022).

55. D. Samigullin, N. Fatikhov, E. Khaziev, A. Skorinkin, E. Nikolsky, E. Bukharaeva, Estimation of presynaptic calcium currents and endogenous calcium buffers at the frog neuromuscular junction with two different calcium fluorescent dyes. Frontiers in Synaptic Neuroscience **Volume** 6-2014 (2015).

56. A. D. Radant, D. J. Dobie, E. R. Peskind, M. M. Murburg, E. C. Petrie, E. D. Kanter, M. A. Raskind, C. W. Wilkinson, Adrenocortical responsiveness to infusions of physiological doses of ACTH is not altered in posttraumatic stress disorder. Frontiers in Behavioral Neuroscience **Volume** 3-2009 (2009).

57. J. M. Waterman, T. M. Cofer, O. M. Von Laue, P. Mateo, L. Wang, M. Erb, Leaf Size Determines Damage- and Herbivore-Induced Volatile Emissions in Maize. Plant, Cell & Environment 10.1111/pce.15355 (2025).

58. T. T. S. Lew, V. B. Koman, K. S. Silmore, J. S. Seo, P. Gordiichuk, S.-Y. Kwak, M. Park, M. C.-Y. Ang, D. T. Khong, M. A. Lee, M. B. Chan-Park, N.-H. Chua, M. S. Strano, Real-time detection of wound-induced H2O2 signalling waves in plants with optical nanosensors. Nature Plants 6, 404–415 (2020).

59. A. D. Steinbrenner, M. Muñoz-Amatriaín, A. F. Chaparro, J. M. Aguilar-Venegas, S. Lo, S. Okuda, G. Glauser, J. Dongiovanni, D. Shi, M. Hall, D. Crubaugh, N. Holton, C. Zipfel, R. Abagyan, T. C. J. Turlings, T. J. Close, A. Huffaker, E. A. Schmelz, A receptor-like protein mediates plant immune responses to herbivore-associated molecular patterns. Proceedings of the National Academy of Sciences 117, 31510–31518 (2020).

60. L. Wang, E. Einig, M. Almeida-Trapp, M. Albert, J. Fliegmann, A. Mithöfer, H. Kalbacher, G. Felix, The systemin receptor SYR1 enhances resistance of tomato against herbivorous insects. Nature Plants 4, 152–156 (2018).

61. J. M. Waterman, C. R. Hall, M. Mikhael, C. I. Cazzonelli, S. E. Hartley, S. N. Johnson, Short-term resistance that persists: Rapidly induced silicon anti-herbivore defence affects carbon-based plant defences. Functional Ecology 35, 82–92 (2021).

62. K. C. Sridhar, N. Hersch, G. Dreissen, R. Merkel, B. Hoffmann, Calcium mediated functional interplay between myocardial cells upon laser-induced single-cell injury: an in vitro study of cardiac cell death signaling mechanisms. Cell Communication and Signaling 18, 191 (2020).

63. X. Wang, H. Fang, Z. Huang, W. Shang, T. Hou, A. Cheng, H. Cheng, Imaging ROS signaling in cells and animals. Journal of Molecular Medicine 91, 917–927 (2013).

64. T.-O. Kiess, J. Kockskämper, SERCA Activity Controls the Systolic Calcium Increase in the Nucleus of Cardiac Myocytes. Frontiers in Physiology **Volume** 10-2019 (2019).

65. R. Vosoughi, Z. Sadeghi Goghari, A. H. Jafari, Modelling system of two insulin-glucose delays to achieve the dynamics of glucose changes. Journal of Biomedical Physics and Engineering 12, 189–204 (2022).

66. D. Morris, J. Maclean, A. J. Black, Computation of random time-shift distributions for stochastic population models. Journal of Mathematical Biology 89, 33 (2024).

67. Y. Jiang, J. Ye, B. Rasulov, Ü. Niinemets, Role of Stomatal Conductance in Modifying the Dose Response of Stress-Volatile Emissions in Methyl Jasmonate Treated Leaves of Cucumber (Cucumis Sativa). International Journal of Molecular Sciences 21 (2020).

68. P. Virtanen, R. Gommers, T. E. Oliphant, M. Haberland, T. Reddy, D. Cournapeau, E. Burovski, P. Peterson, W. Weckesser, J. Bright, S. J. van der Walt, M. Brett, J. Wilson, K. J. Millman, N. Mayorov, A. R. J. Nelson, E. Jones, R. Kern, E. Larson, C. J. Carey, İ. Polat, Y. Feng, E. W. Moore, J. VanderPlas, D. Laxalde, J. Perktold, R. Cimrman, I. Henriksen, E. A. Quintero, C. R. Harris, A. M. Archibald, A. H. Ribeiro, F. Pedregosa, P. van Mulbregt, A. Vijaykumar, A. P. Bardelli, A. Rothberg, A. Hilboll, A. Kloeckner, A. Scopatz, A. Lee, A. Rokem, C. N. Woods, C. Fulton, C. Masson, C. Häggström, C. Fitzgerald, D. A. Nicholson, D. R. Hagen, D. V. Pasechnik, E. Olivetti, E. Martin, E. Wieser, F. Silva, F. Lenders, F. Wilhelm, G. Young, G. A. Price, G.-L. Ingold, G. E. Allen, G. R. Lee, H. Audren, I. Probst, J. P. Dietrich, J. Silterra, J. T. Webber, J. Slavič, J. Nothman, J. Buchner, J. Kulick, J. L. Schönberger, J. V. de Miranda Cardoso, J. Reimer, J. Harrington, J. L. C. Rodríguez, J. Nunez-Iglesias, J. Kuczynski, K. Tritz, M. Thoma, M. Newville, M. Kümmerer, M. Bolingbroke, M. Tartre, M. Pak, N. J. Smith, N. Nowaczyk, N. Shebanov, O. Pavlyk, P. A. Brodtkorb, P. Lee, R. T. McGibbon, R. Feldbauer, S. Lewis, S. Tygier, S. Sievert, S. Vigna, S. Peterson, S. More, T. Pudlik, T. Oshima, T. J. Pingel, T. P. Robitaille, T. Spura, T. R. Jones, T. Cera, T. Leslie, T. Zito, T. Krauss, U. Upadhyay, Y. O. Halchenko, Y. Vázquez-Baeza, SciPy 1.0 Contributors, SciPy 1.0: fundamental algorithms for scientific computing in Python. Nature Methods 17, 261–272 (2020).

69. M. Abramowitz, Stegun, I.A., Handbook of Mathematical Functions with Formulas, Graphs and Mathematical Tables (Dover, New York, ed. 9).

70. A. LoPrete, J. Burge, Bandwidth of Gamma-Distribution-Shaped Functions via Lambert W Function (2025). https://arxiv.org/abs/2509.19307.

71. C. Chen, H. Chen, S. Huang, T. Jiang, C. Wang, Z. Tao, C. He, Q. Tang, P. Li, Volatile DMNT directly protects plants against Plutella xylostella by disrupting the peritrophic matrix barrier in insect midgut. eLife 10, e63938 (2021).

72. G. M. Gurr, J. Liu, J. A. Pickett, P. C. Stevenson, Review of the chemical ecology of homoterpenes in arthropod–plant interactions. Austral Entomology 62, 3–14 (2023).

73. E. Hatano, A. M. Saveer, F. Borrero-Echeverry, M. Strauch, A. Zakir, M. Bengtsson, R. Ignell, P. Anderson, P. G. Becher, P. Witzgall, T. Dekker, A herbivore-induced plant volatile interferes with host plant and mate location in moths through suppression of olfactory signalling pathways. BMC Biology 13, 75 (2015).

74. J. Hua, Modulation of plant immunity by light, circadian rhythm, and temperature. Current Opinion in Plant Biology 16, 406–413 (2013).

75. N. Guayazán Palacios, T. Imaizumi, A. D. Steinbrenner, The Circadian Clock Regulates Receptor-Mediated Immune Responses to a Herbivore-Associated Molecular Pattern. Plant, Cell & Environment **n/a** (2025).

76. J. M. Waterman, C. I. Cazzonelli, S. E. Hartley, S. N. Johnson, Simulated Herbivory: The Key to Disentangling Plant Defence Responses. Trends in Ecology & Evolution 34, 447–458 (2019).

77. T. Degen, C. Dillmann, F. Marion-Poll, T. C. J. Turlings, High Genetic Variability of Herbivore-Induced Volatile Emission within a Broad Range of Maize Inbred Lines. Plant Physiology 135, 1928–1938 (2004).

78. Y. Joo, J. K. Goldberg, L. T. S. Chrétien, S.-G. Kim, I. T. Baldwin, M. C. Schuman, The circadian clock contributes to diurnal patterns of plant indirect defense in nature. Journal of Integrative Plant Biology 61, 924–928 (2019).

79. N. G. Palacios, P. Grof-Tisza, B. Behnken, C. M. Arce, D. Wu, A. F. Chaparro, E. A. Schmelz, T. C. J. Turlings, B. Benrey, A. D. Steinbrenner, A plant immune receptor mediates tritrophic interactions by linking caterpillar detection to predator recruitment. bioRxiv, 2025.07.29.667524 (2025).

80. J. van den Berg, C. Britz, H. du Plessis, Maize Yield Response to Chemical Control of Spodoptera frugiperda at Different Plant Growth Stages in South Africa. Agriculture 11 (2021).

81. D. Maag, C. Dalvit, D. Thevenet, A. Köhler, F. C. Wouters, D. G. Vassão, J. Gershenzon, J.-L. Wolfender, T. C. J. Turlings, M. Erb, G. Glauser, 3-β-d-Glucopyranosyl-6-methoxy-2-benzoxazolinone (MBOA-N-Glc) is an insect detoxification product of maize 1,4-benzoxazin-3-ones. Phytochemistry 102, 97–105 (2014).

82. W. Feller, An Introduction to Probability Theory and Its Applications (Wiley, ed. 2, 1971)vol. 2.

83. N. L. Johnson, S. Kotz, N. Balakrishnan, Continuous Univariate Distributions (Wiley, ed. 2, 1994)vol. 1.

84. D. Kraft, A Software Package for Sequential Quadratic Programming (Wiss. Berichtswesen d. DFVLR, 1988; https://books.google.ie/books?id=4rKaGwAACAAJ)*Deutsche Forschungs-und Versuchsanstalt für Luft-und Raumfahrt Köln: Forschungsbericht*.

85. A. Gelman, J. B. Carlin, H. S. Stern, D. B. Dunson, A. Vehtari, D. B. Rubin, Bayesian Data Analysis (CRC Press, 2013; https://books.google.ie/books?id=eSHSBQAAQBAJ)*Chapman & Hall/CRC Texts in Statistical Science*.

86. G. H. Golub, C. Reinsch, Singular value decomposition and least squares solutions. Numerische Mathematik 14, 403–420 (1970).

87. C. R. Harris, K. J. Millman, S. J. van der Walt, R. Gommers, P. Virtanen, D. Cournapeau, E. Wieser, J. Taylor, S. Berg, N. J. Smith, R. Kern, M. Picus, S. Hoyer, M. H. van Kerkwijk, M. Brett, A. Haldane, J. F. del Río, M. Wiebe, P. Peterson, P. Gérard-Marchant, K. Sheppard, T. Reddy, W. Weckesser, H. Abbasi, C. Gohlke, T. E. Oliphant, Array programming with NumPy. Nature 585, 357–362 (2020).

88. The Pandas Development Team, pandas-dev/pandas: Pandas, version v2.2.2, Zenodo (2024); 10.5281/zenodo.10957263.

89. G. Van Rossum, F. L. Drake, Python 3 Reference Manual (CreateSpace, Scotts Valley, CA, 2009).

90. R Core Team, R: A Language and Environment for Statistical Computing., version 4.4.2, R Foundation for Statistical Computing (2024); https://www.R-project.org/.

91. H. White, A Heteroskedasticity-Consistent Covariance Matrix Estimator and a Direct Test for Heteroskedasticity. Econometrica 48, 817–838 (1980).

92. A. Rohatgi, WebPlotDigitizer, version 5.2 (2025); https://automeris.io.

